# Clostridium difficile exploits a host metabolite produced during toxin-mediated infection

**DOI:** 10.1101/2021.01.14.426744

**Authors:** Kali M. Pruss, Justin L. Sonnenburg

## Abstract

Several enteric pathogens can gain specific metabolic advantages over other members of the microbiota by inducing host pathology and inflammation. The pathogen *Clostridium difficile* (*Cd*) is responsible for a toxin-mediated colitis that causes 15,000 deaths in the U.S. yearly^1^, yet the molecular mechanisms by which *Cd* benefits from toxin-induced colitis remain understudied. Up to 21% of healthy adults are asymptomatic carriers of toxigenic *Cd*^2^, indicating that *Cd* can persist as part of a healthy microbiota; antibiotic-induced perturbation of the gut ecosystem is associated with transition to toxin-mediated disease. To understand how *Cd* metabolism adapts from a healthy gut to the inflamed conditions its toxins induce, we used RNA-seq to define the metabolic state of wild-type *Cd* versus an isogenic mutant lacking toxins in a mouse model. Combining bacterial and mouse genetics, we demonstrate that *Cd* utilizes sorbitol derived from both diet and host. Host-derived sorbitol is produced by the enzyme aldose reductase, which is expressed by diverse immune cells and is upregulated during inflammation, including during *Cd* toxin-mediated disease. This work highlights a mechanism by which *Cd* can utilize a host-derived nutrient generated during toxin-induced disease by an enzyme not previously associated with infection.

---

The enteric pathogen *Clostridium difficile* (*Cd*) causes over 450,000 infections in the United States alone^1^. Symptomatic *Cd* infection (CDI) is mediated by the action of two large glycosylating toxins, TcdA and TcdB, which inactivate the Rho family of GTPases. Importantly, a large proportion of healthy infants and adults are asymptomatic carriers of toxigenic *Cd* (up to 90% and 21% of these populations, respectively^3,4^). It has previously been shown that naturally occurring toxin-negative isolates of *Cd* fail to cause disease and elicit a metabolic profile distinct from toxigenic strains^5^. Markers of inflammation are a better predictor of CDI severity than pathogen burden^6^: the host immune response plays a crucial role in dictating the status (symptomatic or asymptomatic) of CDI, but how *Cd* may exploit the distinct ecological niches of a healthy versus inflamed gut remains unknown^7^.

Many enteric pathogens cause disease in the gut by expressing virulence factors to create and maintain a modified niche that provides a competitive advantage over other community members^7,8^: *Salmonella enterica* serovar Typhimurium^9,10^, *Citrobacter rodentium*^11^, enterohemorrhagic *Escherichia coli* (EHEC)^12,13^, and *Vibrio cholerae*^14^ couple metabolic strategies to specific aspects of host response. This foundation of knowledge for how pathogens benefit from the pathology they induce provides a useful template for pursuing an understudied aspect of *Cd* pathogenesis. We hypothesized that toxin production alters the biochemical landscape of the gut and that *Cd* alters its metabolism accordingly. To understand the influence of toxin-mediated inflammation on *Cd* metabolism, we utilized gnotobiotic mice with wild-type *Cd* or an isogenic toxin-deficient mutant^15^. RNA-seq allowed identification of metabolic pathways in *Cd* that were specific to inflammatory or non-inflammatory conditions. Using these controlled diet and gnotobiotic conditions, we predicted that toxin-dependent shifts in the metabolic behavior of *Cd* would reflect alterations in nutrient availability in the lumen due to host-derived metabolites.

## Toxin-induced inflammation leads to extensive differential *Cd* gene expression *in vivo*

Toxin production conferred an advantage in relative abundance to WT *Cd* in the presence of a commensal community (**Extended Data Fig. 1a**). To investigate toxin-dependent changes to *Cd* metabolism, we mono-colonized germ-free mice with wild-type *Cd* 630ΔErm or *Cd* 630ΔErmTcdA^-^B^-^, an isogenic mutant that lacks the ability to produce toxins **(Extended Data Fig. 1b)**^15^. Colonization with the toxin-deficient mutant (here referred to as Tox-) resulted in negligible inflammation in the proximal colon of mice compared to the wild-type (WT) strain (**Fig. 1a**). Histopathological scoring confirmed that mice infected with WT *Cd* had higher numbers of inflammatory cell infiltrates, mucosal hyperplasia, and vascular congestion (**Extended Data Table 1**).

**Fig. 1.**
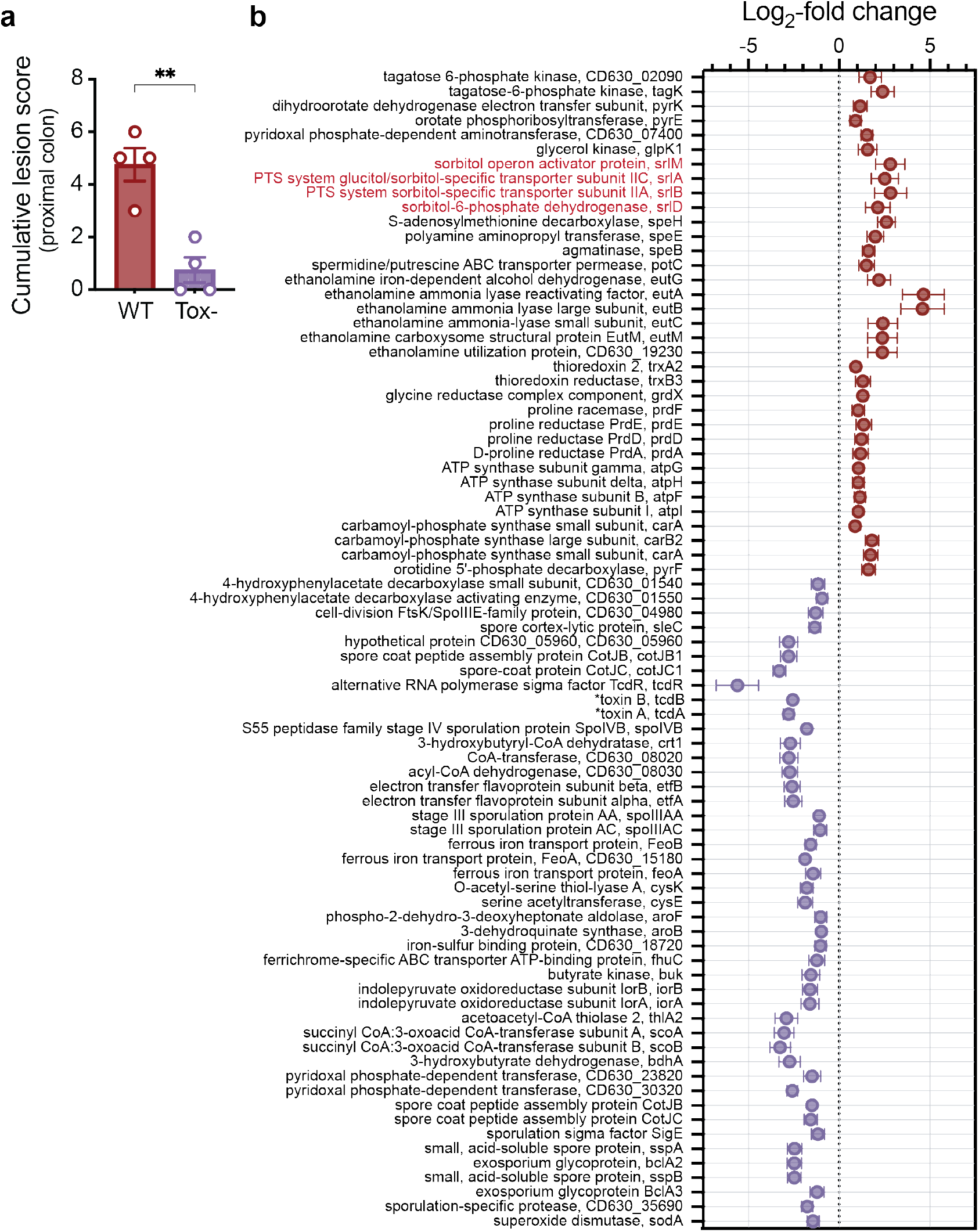
Toxin-induced inflammation leads to extensive differential gene expression of *Cd in vivo*. **a**, Blinded histopathological scoring of the proximal colon for mice infected with WT and Tox-mutant *Cd* at day 3 post-colonization (mean +/-SEM, ** *P* = 0.0023, two-tailed unpaired Student’s t test). **b**, Fold-change (mean +/-SEM, n=3-4/group) of a subset of *Cd* genes significantly differentially expressed between WT and Tox-*Cd* at day 3 post-colonization in cecal contents of gnotobiotic mice. The two toxin proteins are denoted by asterisks and the putative sorbitol utilization locus is highlighted in red. Genes included were involved in iron transport, sporulation, virulence, and metabolism (for metabolic genes, >= 2 genes in an operon).

After 3 days of infection, we isolated total RNA from cecal contents for RNA-seq. 331 genes were differentially expressed between wild-type and toxin-deficient *Cd* (adjusted *P*-value < 0.05, Wald test with Benjamini-Hochberg correction). Several metabolic pathways were enriched due to the presence or absence of toxin *in vivo* (**Extended Data Figs. 1c,d**). Genes involved in sporulation, conjugative transposons, iron transport and butyrate production were more highly expressed in the absence of toxin-induced inflammation (**Extended Data Table 2**). Interestingly, more reads mapped to the mutated toxin genes themselves in Tox-(**Fig. 1b**), despite the strain’s inability to produce toxin proteins. Genes involved in the metabolism of ethanolamine, a host-derived molecule utilized by other enteric pathogens within the gut environment^16–18^, were more highly expressed in WT (**Fig. 1b**).

## Molecular basis for *Cd* sorbitol metabolism

An operon putatively annotated for sorbitol utilization was more highly expressed in WT than in Tox-*Cd* (**Fig. 1b,2a**). Sorbitol metabolism has been long described in *Cd*^19^ but not investigated *in vivo*. Based on previous work in *E. coli*^20,21^ and *Streptococci*^22^, it is predicted that sorbitol enters the cell via a specific phosphotranferase system (PTS), generating a 6-phospho derivative. Sorbitol-6-phosphate dehydrogenase acts on D-sorbitol-6P, oxidizing its 2-alcohol group to a keto group, using NAD^+^ as a cofactor. The resulting keto sugar, ß-D-fructofuranose-6P is an intermediate of glycolysis (**Extended Data Fig. 2a**). Both WT and Tox-*Cd* were able to grow using sorbitol as a sole carbohydrate source *in vitro* (**Fig. 2b, Extended Data Fig. 2b**), indicating that lower expression of the sorbitol utilization operon in Tox-*in vivo* was not due to an inability to utilize the substrate.

**Fig. 2.**
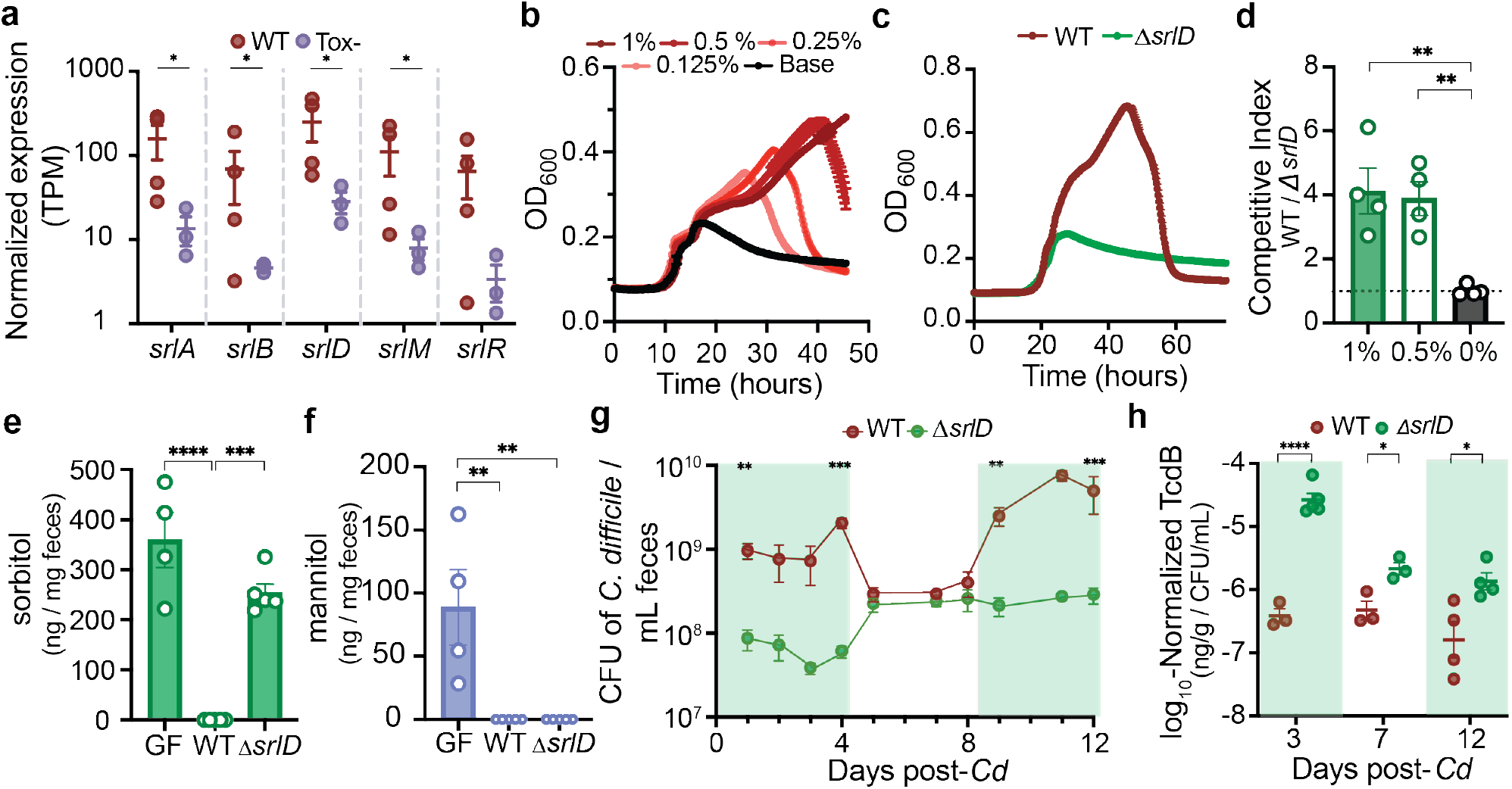
A putative sorbitol utilization locus is responsible for *Cd* metabolism of sorbitol *in vitro* and *in vivo*. **a**, Normalized expression values (Transcripts per Million) for genes involved in the putative sorbitol utilization locus. **b**, *Cd* dose-dependent growth in minimal medium supplemented with sorbitol (mean +/-SEM shown, n=5 replicates per condition). **c**, A *Cd* sorbitol dehydrogenase mutant (*ΔsrlD*) is unable to achieve increased growth yield in minimal medium supplemented with 0.5% w/v sorbitol (mean +/-SEM, n=5 replicates per condition. **d**, WT *Cd* outcompetes *ΔsrlD in vitro* after 24 hours when sorbitol (0.5%, 1% w/v) is added to minimal medium but not when sorbitol is absent (0%, mean +/-SEM, one-way ANOVA F_(2,9)_=11.37 with Dunnett’s multiple comparisons using 0% as control group). **e**, Colonization of mice fed standard diet with *ΔsrlD Cd* (*ΔsrlD*) does not alter fecal sorbitol levels compared to germ-free mice (GF), whereas colonization with WT *Cd* (WT) reduces sorbitol levels below the limit of detection (mean +/-SEM, sorbitol: F_(2,11)_=40.25, one-way ANOVA with Tukey’s multiple comparisons). **f**, Mannitol in GF mouse feces is depleted upon colonization with either WT or *ΔsrlD Cd* (F_(2,11)_=11.56, one-way ANOVA with Tukey’s multiple comparisons). **g**, Exogenously-provided sorbitol increases abundance of WT *Cd in vivo* but not the *ΔsrlD* mutant. Green boxes indicate days for which sorbitol was supplemented in drinking water (1% w/v, mean +/-SEM, n=3-5, multiple t-tests with Holm-Sidak method for multiple comparisons). **h**, Normalized TcdB abundance in feces of mice infected with either WT or *ΔsrlD Cd*. The inability to utilize sorbitol leads to increased toxin production *in vivo* (mean +/-SEM, multiple t-tests with Holm-Sidak method for multiple comparisons). For all panels, * *P* < 0.05, ** *P* < 0.01, *** *P* < 0.001, **** *P* < 0.0001.

To determine whether the putative sorbitol locus was responsible for demonstrated growth on sorbitol (**Fig. 2b**), we created a deletion mutant of the sorbitol dehydrogenase gene (Δ*srlD*). The deletion mutant was unable to achieve increased growth on minimal medium supplemented with sorbitol (**Fig. 2c, Extended Data Fig. 2c**) despite maintaining induction of the *srl* locus (**Extended Data Fig. 2d**, genes annotated as sorbitol dehydrogenase, PTS transporter subunit, and transcription anti-terminator respectively). Furthermore, WT outcompeted the Δ*srlD* mutant *in vitro* only when sorbitol was added to the medium (**Fig. 2d**).

## *Cd* consumes sorbitol *in vivo*

Next, we investigated sorbitol utilization by *Cd* within the mouse gut. The addition of exogenous sorbitol in drinking water led to up-regulation of the *Cd srlD* gene (**Extended Data Fig. 2e**), indicating that increased availability of sorbitol led to induction of the utilization locus. It was previously shown in an untargeted metabolomic screen that, after a course of antibiotics rendering mice susceptible to CDI, sorbitol and mannitol were the most enriched metabolites relative to pre-or post-antibiotic recovery states^23^. We developed a GC-MS-based mass spectrometry assay to differentiate and quantify both sorbitol and mannitol and demonstrated that our antibiotic pre-treatment model for *Cd* infection in conventional mice led to an increase in sorbitol and mannitol in stool (**Extended Data Fig. 2f**).

Sorbitol and its stereo-isomer mannitol are by-products of photosynthesis and serve as carbon storage molecules in plants^24^; we predicted that both would be abundant in standard mouse chow. Accordingly, germ-free mice exhibited high levels of free sorbitol and mannitol in feces (**Fig. 2e,f**). Subsequent colonization of germ-free mice with WT *Cd* led to complete consumption of both sugar alcohols, while colonization with the Δ*srlD* strain did not alter levels of sorbitol (**Fig. 2e**), but did deplete mannitol, indicating that *srlD* is highly specific to sorbitol (**Fig. 2f**). The inability to utilize sorbitol led to attenuated colonization 24 hours after infection in conventional mice fed standard diet (**Extended data Fig. 2g**). The lower colonization level of the *ΔsrlD* mutant was accompanied by increased relative production of toxin (**Extended data Fig. 2h**), consistent with toxin production by *Cd* in nutrient-limited and stressed conditions^7^.

To determine whether sorbitol availability influenced *Cd* abundance in the gut, we placed gnotobiotic mice harboring a defined community of bacteria on a fully-defined diet devoid of sorbitol, and subsequently infected with either WT *Cd* or *ΔsrlD*. When sorbitol was supplemented in drinking water (1% w/v), WT reached an order of magnitude higher density than *ΔsrlD*; when sorbitol was removed from drinking water, the abundance of WT dropped to levels indistinguishable from *ΔsrlD* (**Fig. 2g**). As seen in conventional mice, the inability to utilize sorbitol resulted in increased toxin production on a per-cell basis (**Fig. 2h**).

We tested whether exogenous mannitol supplementation would lead to an increase in *Cd* abundance as seen with sorbitol (**Fig. 2g**). 1% sorbitol or 1% mannitol supplementation in drinking water to gnotobiotc mice harboring a defined consortium of bacteria led to an increased density of WT *Cd* (days 0-6 and 10-14) compared to when no sorbitol or mannitol was provided (days 7-10, **Extended data Fig. 3a**). 1% mannitol in drinking water allowed the *ΔsrlD* mutant to achieve increased abundance (**Extended data Fig. 3b**) and incite an equivalent histopathology score as WT (**Extended data Figs. 3c,d**), demonstrating that provision of a sugar alcohol that is stereoisomeric to sorbitol enabled mutant outgrowth, whereas sorbitol did not.

## Absence of sorbitol metabolism leads to increased *Cd* toxin production *in vitro* and *in vivo*

We sought to define the influence of sorbitol metabolism on toxin production in *Cd*. 0.5% or 1% sorbitol supplementation (w/v) in minimal media led to lower expression of *tcdA* and *tcdC*, the holin-like protein required for toxin secretion^26,27^, compared to un-supplemented base medium (**Extended data Fig. 4a**). *In vitro* culture experiments are informative for how specific nutrient conditions impact *Cd* toxin expression; toxin production within the host gut (*in vivo*) represents a complex integration of many cues in addition to nutrients, such as host inflammatory mediators and metabolites from other microbes. Provision of exogenous sorbitol or mannitol *in vivo* (in drinking water) led to lower levels of toxin production by WT *Cd* compared to when no sorbitol or mannitol was supplemented (**Extended data Fig. 4b**). In contrast, only mannitol, not sorbitol, decreased toxin expression in the *ΔsrlD* mutant strain of *Cd* that is unable to use sorbitol (**Extended data Fig. 4c**). Our data support a model in which amplified toxin production occurs in the absence of *Cd* sorbitol metabolism, potentially as a mechanism to procure the sugar alcohol. It is important to note that host inflammation during CDI is a function of cumulative toxin amount and several factors including changes in host immune response, gut motility, and dynamics of the resident microbiota, including the changing density and biogeography of *Cd*.

When our data is compared with previous reports examining the impact of simple sugars on different aspects of *Cd* lifestyle (host colonization, sporulation, toxin expression) there appear to be important and complex differences that likely reflect the individual history and source of each nutrient. Recent data demonstrated that a dietary sugar additive to which some epidemic *Cd* strains recently adapted, trehalose, led to induction of *Cd* virulence genes^28^. Administration of the disaccharide trehalose to mice led to decreased host survival during CDI; a *Cd* mutant unable to metabolize trehalose exhibited lower toxin production and increased survival in a mouse model^25^. A recent study of 906 whole-genome *Cd* isolates showed that transporters and metabolic pathways for many sugars, including glucose, mannitol, sorbitol, and fructose (also a product of sorbitol metabolism) are under positive selection in many *Cd* strains^26^. In these studies, administration of glucose and fructose, but not ribose (for which metabolism is not positively selected), was shown to enhance *Cd* sporulation and gut colonization in mice. The sources of these sugars vary: some have been long-term components of the human diet, others recent additions as dietary additives, and a subset also produced by the host. Each has undergone a unique rate of increase in the industrialized diet^27^, and their impacts on *Cd* physiology and evolution is an important area of investigation.

## Host aldose reductase is an immune cell-associated gene that responds to *Cd* infection

In our initial gnotobiotic RNA-seq conditions investigating inflammation-dependent changes to *Cd* metabolism (**Fig. 1b**), mice were fed identical diets. We thus hypothesized that sorbitol utilization during toxin-dependent disease may be due to host production of sorbitol during an inflammatory response. In the mammalian polyol pathway, the enzyme aldose reductase (AR) reduces glucose to sorbitol; sorbitol dehydrogenase subsequently oxidizes sorbitol to fructose (**Fig. 3a**), which is a substrate for glycolysis. Under hyperglycemic conditions, up to 30% of glucose is shunted through the polyol pathway^28^.

**Fig. 3.**
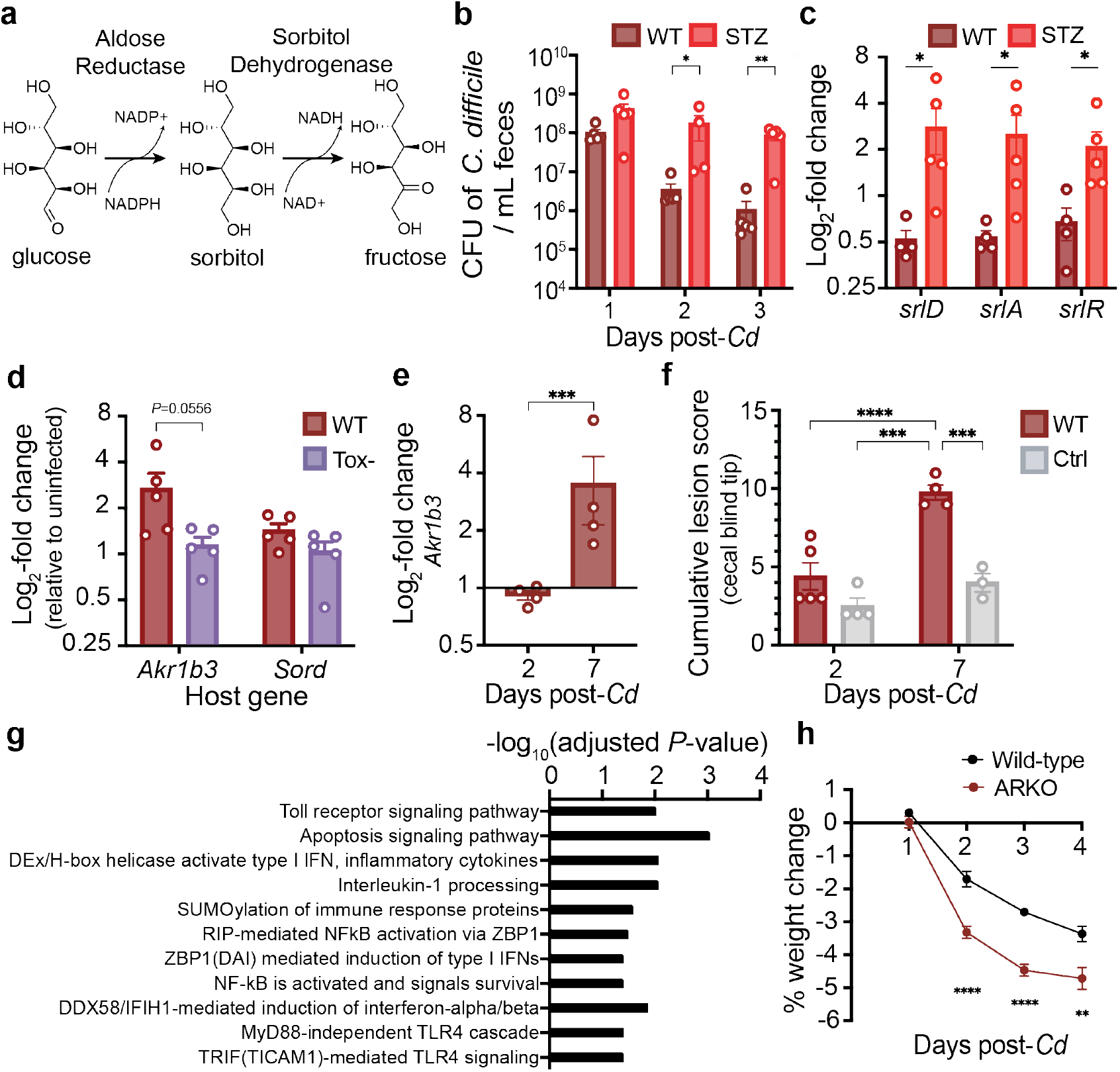
Host aldose reductase responds to *Cd* infection. **a**, Schematic of the host polyol pathway. Aldose reductase reduces glucose to sorbitol using NADPH as a cofactor. Sorbitol is oxidized to fructose by sorbitol dehydrogenase, generating NADH. **b**, *Cd* fecal density in conventional mice with or without Streptozotocin treatment (mean +/-SEM, Mann-Whitney). **c**, Relative expression levels of the *Cd* sorbitol utilization locus in Streptozotocin-treated mice versus controls as measured by qRT-PCR of RNA isolated from feces 3 days after infection (mean +/-SEM, Mann-Whitney). **d**, Relative expression levels of AR (*Akr1b3*) and sorbitol dehydrogenase (*Sord*) in the proximal colon of mice infected with hypervirulent WT (R20291) or Tox-(R20291 TcdA^-^TcdB^-^CDT^-^) *Cd* (5 days post-infection, expression levels are normalized to an uninfected control group not shown; Mann-Whitney, bars denote mean + SEM,). **e**, Proximal colonic expression of AR (*Akr1b3*) in conventional mice 2 and 7 days post infection with *Cd*. **f**, Blinded histopathological scoring of conventional mice cecum at 2 and 7 days post infection with *Cd* relative to uninfected controls; all mice treated with 1 mg Clindamycin 24 hours prior to infection (mean +/-SEM, one-way ANOVA F_(3,12)_=20.83 with Tukey’s post-hoc comparisons). **g**, Host pathways that are significantly positively correlated with host aldose reductase expression in response to purified *Cd* TcdA. **h**, Aldose reductase knockout mice lose significantly more weight than wild-type mice when infected with wild-type *Cd* (n=15 wild-type mice and n=16 ARKO mice from two independent experiments; mean +/-SEM, mixed-effects analysis with multiple comparisons.). For all panels, * *P* < 0.05, ** *P* < 0.01, *** *P* < 0.001, **** *P* < 0.0001. In **c** and **e**, one outlier was tested for and removed with robust nonlinear regression, Q = 0.2%; original data is provided in Extended Data Figs. 5c and 6f.

It has previously been shown that increasing blood glucose leads to higher activity of AR^29,30^. Streptozotocin, a glucosamine-nitrosourea compound recognized by GLUT2 transporters, is toxic to the insulin-producing beta cells of the pancreas. We utilized a streptozotocin-induced model of hyperglycemia in conventional (**Extended Data Fig. 5a**) or germ-free mice (**Extended Data Fig. 5b**). STZ-treated conventional mice were more susceptible to *Cd* colonization (**Fig. 3b**) and *Cd* upregulated the locus for sorbitol utilization during hyperglycemia (**Fig. 3c, Extended data Fig. 5c)** independent of a change to toxin production (**Extended Data Fig. 5d**), indicating increased availability of sorbitol from host tissue. As sorbitol and mannitol are both components of diet, but only sorbitol can be generated by the mammalian host, we predicted that the two sugar alcohols would differentially affect *Cd. In vitro* transcriptional profiling confirmed^31^ distinct metabolic programs and physiological responses by *Cd* to sorbitol versus mannitol (see Supplementary Text). Intriguingly, there is a higher incidence of *Cd* in diabetic patients^32,33^, although the extent to which host AR activity and sorbitol utilization by *Cd* contributes to this is currently unknown.

While the polyol pathway has primarily been studied in diabetes, there is some evidence suggesting a role for AR in the innate immune response. AR inhibition prevents expression of various inflammatory markers: cytokines, including TNF, IL-6, IL-1, IFNg; chemokines, such as MCP-1 and MIP-1; additional inflammatory proteins, including iNOS and Cox-2^34^. To identify the principal host tissues that express AR, we examined published single-cell RNA-seq data from the *Tabula muris* study, which performed single-cell RNA-seq on 20 mouse organs^35^. AR (*Akr1b3*) is most highly expressed in several bone marrow-derived cell types, including megakaryocyte-erythroid progenitors, granulocyte monocyte progenitors, pre- and natural killer cells (also in liver), multipotent progenitor cells, common lymphoid progenitors and basophils (**Extended Data Fig. 6a**)^35^, further indicating a role in immune function.

To determine what cell types express AR within the gastrointestinal tract, we analyzed single cell RNA-seq data from tissues taken from mouse large intestine^36^ and human colonic explants^37^. *Akr1b3* in mice (**Extended data Fig 6b**) and *Akr1b1* (**Extended data Fig. 6c**), the human homolog, were expressed at high prevalence by several immune cells in the gut. Notably, sorbitol dehydrogenase was not comparably expressed in these cells, but was most expressed in enterocytes, goblet cells, and stem cells (**Extended data Fig. 6b,c)**. Several immune cell types increased AR expression (*Akr1b1* or *Akr1b3*) during inflammatory conditions in the gut of mice (during *H. polygyrus* infection)^36^ and humans (inflammatory tissue from ulcerative colitis patients)^37^ relative to uninflamed controls (**Extended data Fig. 6d,e**).

To increase the host inflammatory response during CDI, we infected mice with the hypervirulent *Cd* strain R20291 or its isogenic triple-toxin knockout^38^. In mice infected with WT R20291, AR was more highly expressed in proximal colonic tissue compared to mice infected with Tox-, whereas there was no change in expression to host sorbitol dehydrogenase, indicating that AR selectively responded to *Cd*-induced inflammation (**Fig. 3d**). In conventional mice infected with *Cd* 630ΔErm, AR was more highly expressed in the proximal colon (**Fig. 3e, Extended data Fig. 7f**), concomitant with increased inflammation and tissue damage (**Fig. 3f**). When purified *Cd* TcdA was injected into the ceca of mice, host AR was significantly up-regulated in cells isolated from the epithelial layer of the cecum compared to sham-injected controls (adjusted *P*-values for three *Akr1b3* microarray probes = 0.0175, 0.0198, 0.0256, moderated t-statistic with Benjamini-Hochberg false discovery rate correction)^39^. Thus, host AR responds to three different models of CDI inflammation.

Finally, we conducted a differential correlation analysis for genes that were positively correlated with AR (*Akr1b3*) in response to purified *Cd* TcdA^39^ but negatively or non-correlated in control mice (sham injection) 16 hours post-injection. Pathway enrichment analysis revealed several host immune pathways that were significantly positively correlated with host AR (**Fig. 3g**). Taken together, these data demonstrate that AR is a novel component of the host immune response during infection.

To decrease host AR activity in gnotobiotic mice, we administered the AR inhibitor epalrestat via gavage. Although epalrestat reduced *Cd* density in a mouse model of infection consistent with loss of host sorbitol, after careful follow-up investigation, we determined that the drug acted as a novel antibiotic for *Cd* (see Supplementary Text). Thus, the influence of epalrestat on *Cd in vivo* could be the result of direct inhibitory effects, or indirect effects mediated through inhibition of sorbitol production. As an alternative, we proceeded with genetic ablation of AR, employing AR knockout mice (ARKO) harboring a conventional microbiota. Further supporting the role of AR in the immune response, ARKO mice lost weight more rapidly than wild-type mice over the course of post-antibiotic *Cd* infection (**Fig. 3h**).

## *Cd* utilizes host-derived sorbitol

To determine whether *Cd* utilizes host-derived sorbitol, we co-infected conventional mice with equal amounts of WT *Cd* and the isogenic mutant *ΔsrlD* in the absence of dietary sorbitol. On a fully-defined diet devoid of sorbitol, WT outcompeted the *ΔsrlD* mutant by up to five orders of magnitude (**Fig. 4**, black bars). In mice lacking the ability to generate sorbitol (ARKO mice), wild-type *Cd* lost its competitive advantage over the mutant unable to metabolize sorbitol (**Fig. 4**, red bars). Exogenous administration of sorbitol to ARKO mice on a defined diet restored the competitive advantage to WT *Cd* (**Fig 4**, green bars). Although we cannot exclude the possibility that increased toxin production by *ΔsrlD* contributes to its disadvantage, previous studies in gut bacteria have demonstrated that sustained over-expression of single protein-coding genes does not diminish competitive fitness *in vivo* over weeks^40^. Our data are consistent with *Cd* utilization of sorbitol produced by host AR during infection.

**Fig. 4.**
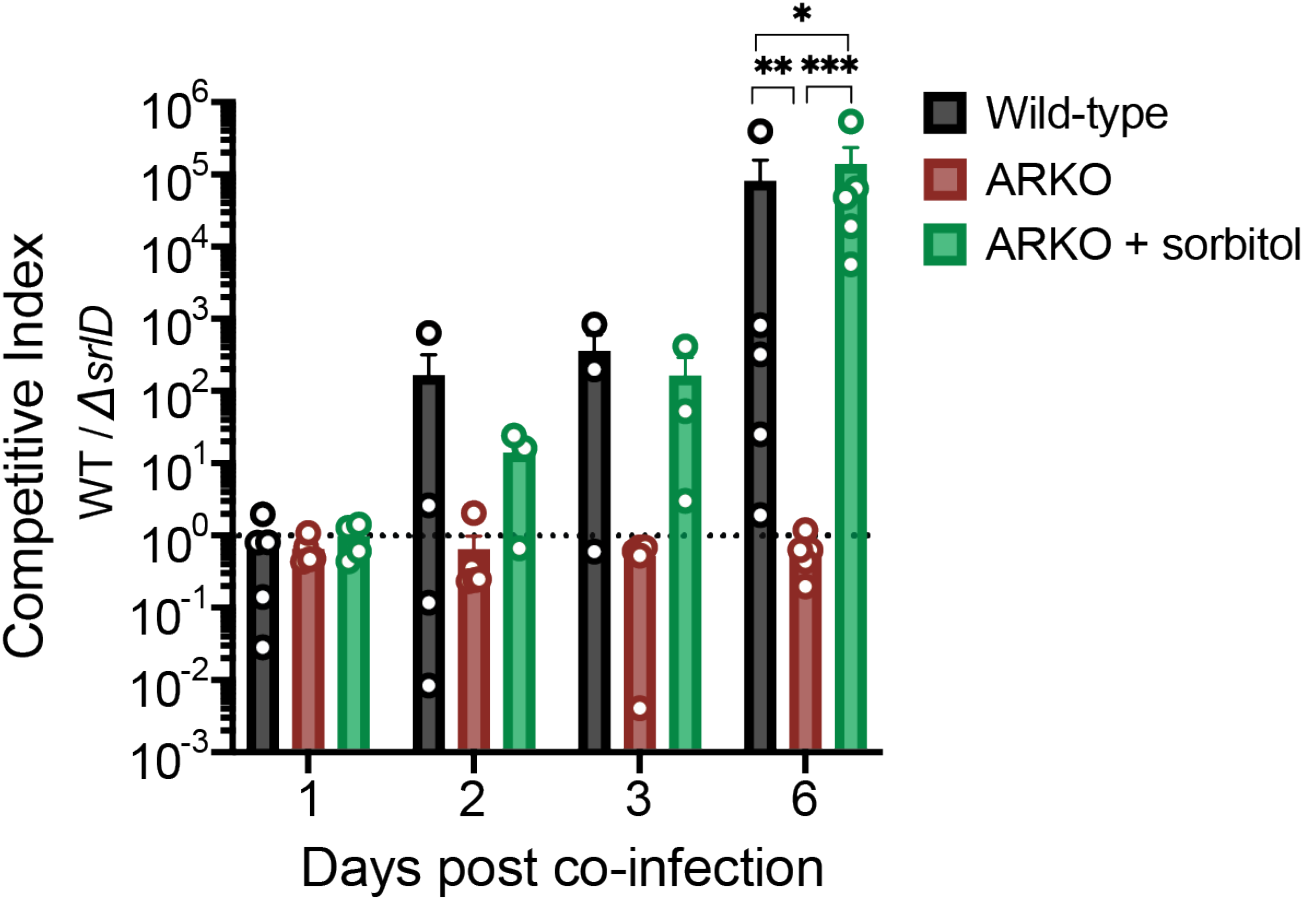
*Cd* utilizes host-derived sorbitol to achieve high densities within the gut. Mice were placed on a fully-defined diet devoid of sorbitol 1 day prior to treatment with Clindamycin. Wild-type *Cd* out-competes the Δ*srlD* mutant in wild-type mice, but not in ARKO mice. Administration of exogenous sorbitol to ARKO mice complements host genetic deficiency and restores the competitive advantage to wild-type *Cd* (mean +/-SEM, n=4-5 mice per group. F_(2,12)_=20.02, one-way ANOVA at day 6 with post-hoc multiple comparison; * *P* < 0.05, ** *P* < 0.01, *** *P* < 0.001). Dashed line at a competitive index value of 1, denoting equal abundance.

Here, we demonstrate for the first time that *Cd* induces and utilizes a host-derived substrate in a mouse model of infection. Our data along with data from previous studies suggest that *Cd* may exist in a commensal state or pathogenic state^2,5,7,41^ (**Extended Data Fig. 7**). Endogenous levels of sorbitol and mannitol are low, as the sugar alcohols are competitively consumed by the commensal microbial community. Numerous studies have shown that upon disturbance of the microbiota, *Cd* can capitalize upon resources that become available during perturbation^8,42,43^, including the transient increase in abundance of freed sorbitol and mannitol (**Extended Data Fig. 2f, ref**.^23^). Subsequently, diminished levels of perturbation-associated nutrients leads to increased toxin production by *Cd*^44^. With toxin-induced inflammation, AR expression increases. The resulting host-produced sorbitol is freed from tissue via cellular damage associated with toxin production and represents a diet-independent resource for the pathogen during inflammation. The trans-kingdom metabolic interactions elucidated here suggest two distinct therapeutic strategies, one targeting *Cd* metabolism relevant to the inflammatory environment it creates, and a second focused on host pathways or metabolites that support *Cd* persistence *in vivo*.

**Extended Data Fig. 1.**
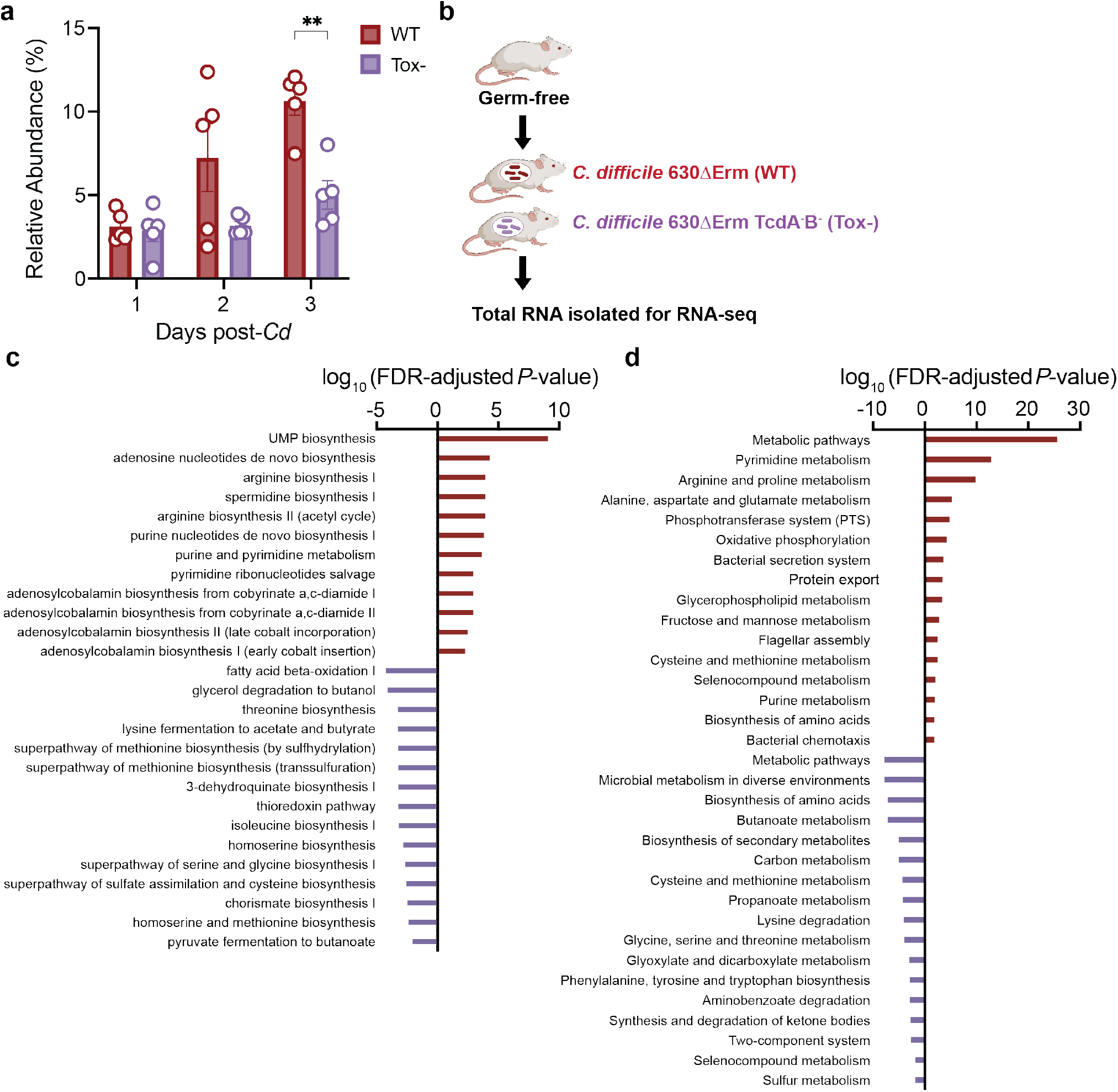
*Cd* toxin production confers an advantage and alters metabolic pathways *in vivo*. **a**, Toxin production (WT) confers an advantage in *Cd* relative abundance in the presence of a defined community (multiple unpaired t-tests with Welch’s correction, two-stage step-up procedure of Benjaini, Krieger, and Yekutieli. ** adjusted *P*-value < 0.01). **b**, Transcriptional profiling experimental design: germ-free mice on standard diet were mono-colonized with either wild-type *Cd* 630ΔErm (WT) or 630ΔErmTcdA-B-(Tox-) (n=4/group). Three days post-infection, total RNA was isolated from cecal contents for RNA-seq. Significantly enriched Ecocyc **(c)** and KEGG **(d)** pathways based on genes differentially expressed during WT (positive, red bars) or Tox-*Cd* (negative, purple) infection (hypergeometric distribution followed with false discovery rate corrected-*P* values).

**Extended Data Fig. 2.**
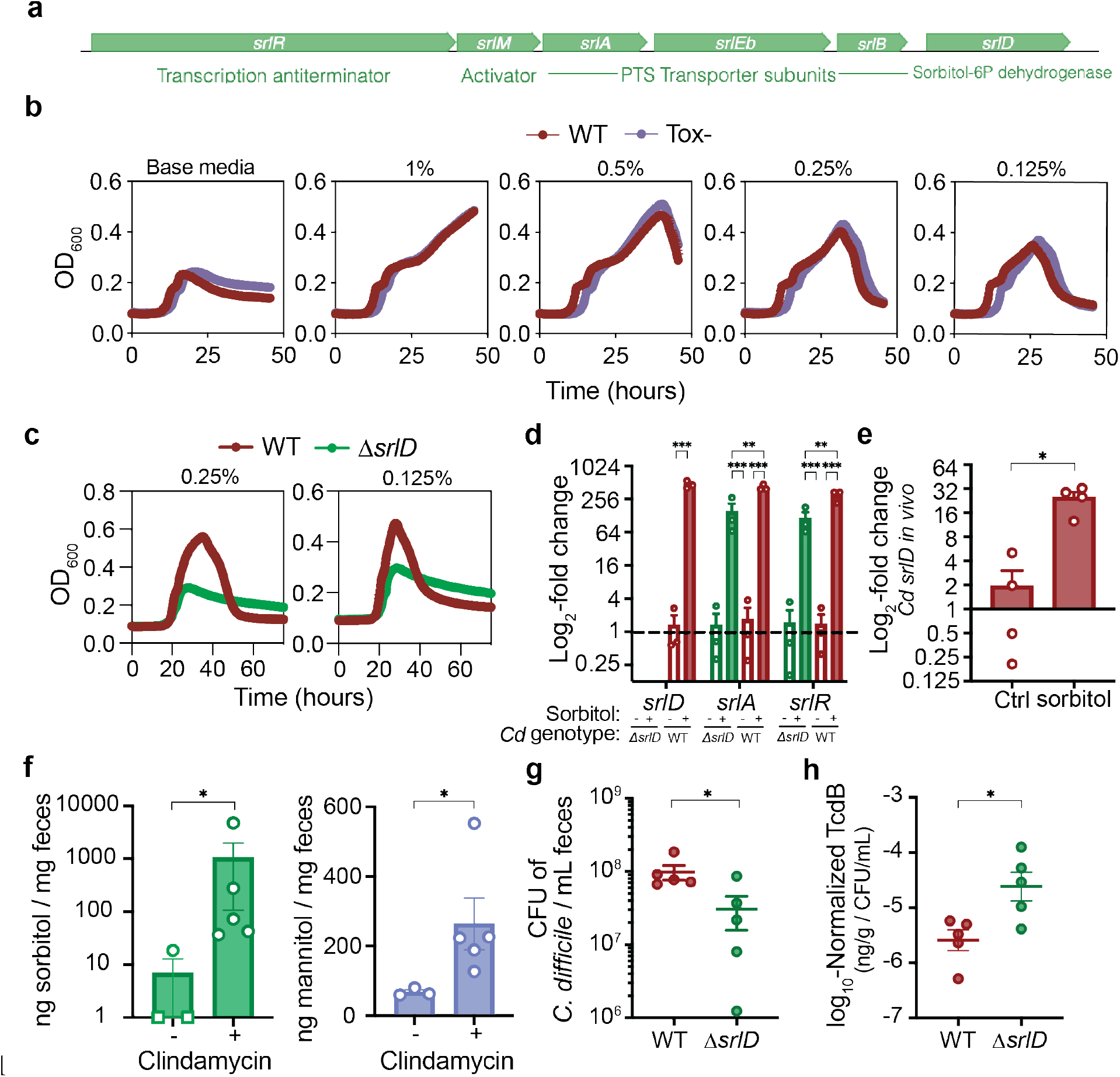
Sorbitol impacts *Cd* growth, gene expression, and increases in the mouse gut after antibiotic treatment. **a**, Schematic overview of the putative sorbitol utilization locus in *Cd* 630ΔErm. The operon consists of three PTS transporter subunits, a 6-phosphate dehydrogenase, an activator and an anti-terminator. **b**, WT (red) and Toxin-deficient (Tox-, purple) *Cd* grow comparably in minimal medium supplemented with various concentrations of sorbitol (mean +/-SEM, n=5 replicates per condition). **c**, *ΔsrlD* mutant is unable to achieve increased growth yield with 0.25% (left) or 0.125% (right) w/v sorbitol supplemented to minimal medium **d**, Addition of sorbitol to minimal medium leads to up-regulation of genes *srlD* (WT), *srlA*, and *srlR* (WT and *ΔsrlD*) compared to base medium (mean +/-SEM, n=3 replicates each condition. Expression levels normalized to WT *Cd* in unsupplemented base medium, dotted line indicates baseline expression of 1; *srlD*: unpaired t-test, *srlA* and *srlR*: one-way ANOVA with post-hoc comparisons; *srlA*: F_(3,8)_ =31.85, *srlR*: F_(3,8)_=27.25.). **e**, Sorbitol administered to mice mono-colonized with wild-type *Cd* 630ΔErm leads to induction of *srlD in vivo* (mean +/-SEM, n=4/group, t-test with Welch’s correction). **f**, 1 mg Clindamycin treatment leads to increased sorbitol and mannitol in stool from conventional mice on standard diet (mean +/-SEM, Mann-Whitney. Sorbitol levels were below the limit of detection for two of three pre-antibiotic treatment samples and are denoted by squares at a value of 1. Samples are combined from 3 independent experiments). **g**, *ΔsrlD Cd* is attenuated in colonization of conventional mice fed a standard diet compared to WT *Cd* (t-test with Welch’s correction). **h**, Toxin B detected by ELISA in fecal pellets of conventional mice 24 hours post-infection with WT or *ΔsrlD Cd*; values were normalized to the absolute abundance of *Cd* from the same stool sample (mean +/-SEM, Mann-Whitney). For all panels, * *P* < 0.05, ** *P* < 0.01, *** *P* < 0.001.

**Extended Data Fig. 3.**
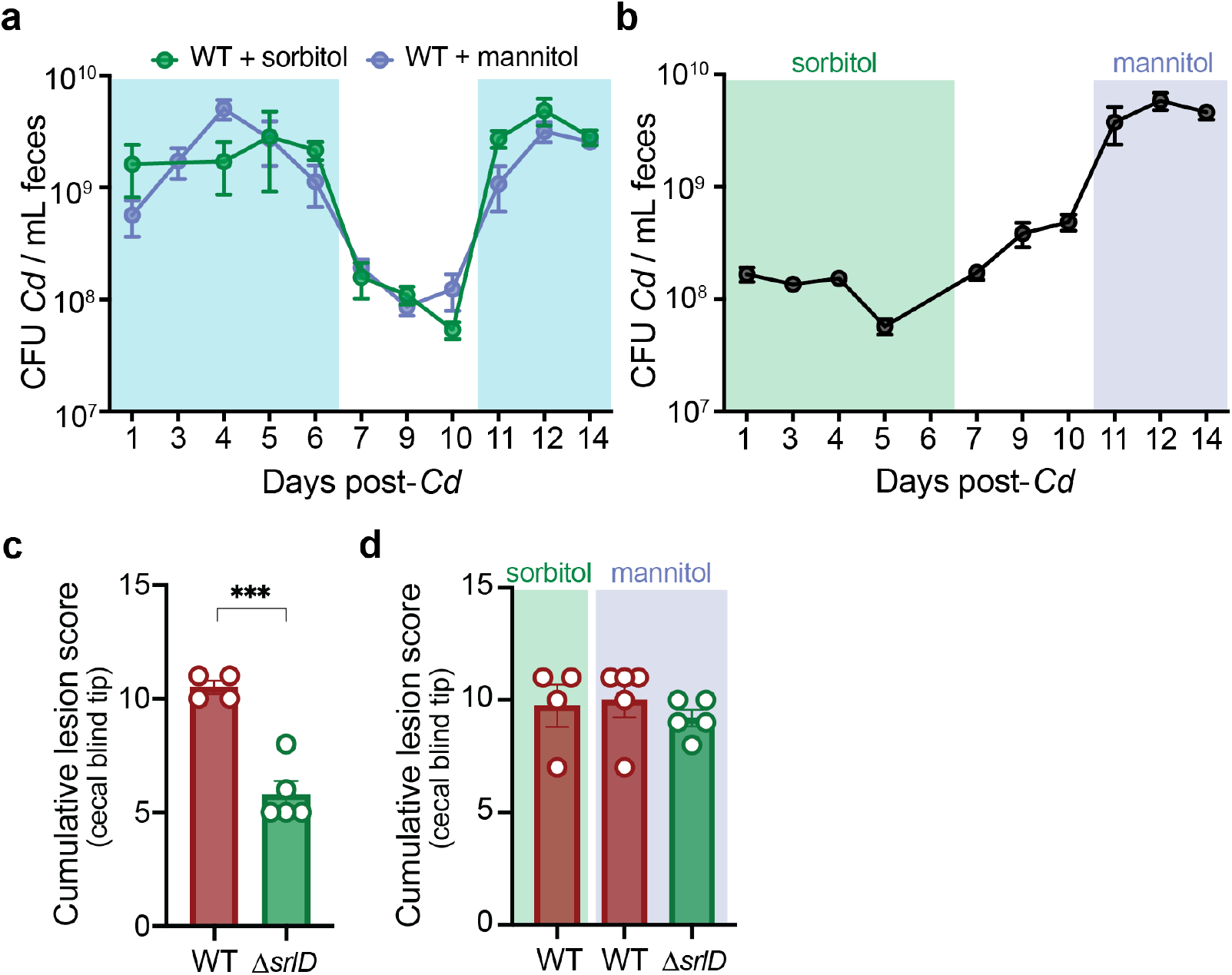
Dietary sorbitol or mannitol availability increases *Cd* density *in vivo*. **a**, 1% sorbitol (green symbols) or 1% mannitol (purple symbols) were provided in drinking water to gnotobiotic mice harboring a defined consortium of bacteria for 6 days (days 0-6). Absolute abundance of WT *Cd* decreases when sorbitol and mannitol are removed from drinking water (days 7-10). Replacing 1% sorbitol and mannitol in drinking water (days 11-14) restores increase in absolute abundance (mean +/-SEM, n=4-5 mice/group; shaded boxes denote duration for which sorbitol or mannitol were provided). **b**, 1% sorbitol was provided in drinking water (days 0 to 6, green box) to mice colonized with a defined community and subsequently infected with *ΔsrlD Cd*. Supplementation of 1% mannitol in drinking water leads to an increase in *ΔsrlD* abundance (days 11-14, purple box) relative to sorbitol supplementation (mean +/-SEM, n=5 mice). **c**, *ΔsrlD Cd* incites a lower histopathological score than WT *Cd* when 1% sorbitol is supplemented in drinking water (12 days post-infection, mean +/-SEM, unpaired Student’s t-test, *** *P* < 0.001). **d**, No significant differences in blinded histopathological scoring at end point (day 14) in the cecal blind tip of mice infected with WT *Cd* when sorbitol or mannitol is supplemented (as in **a**) or when mannitol is supplemented to *ΔsrlD Cd* (**b**).

**Extended Data Fig. 4.**
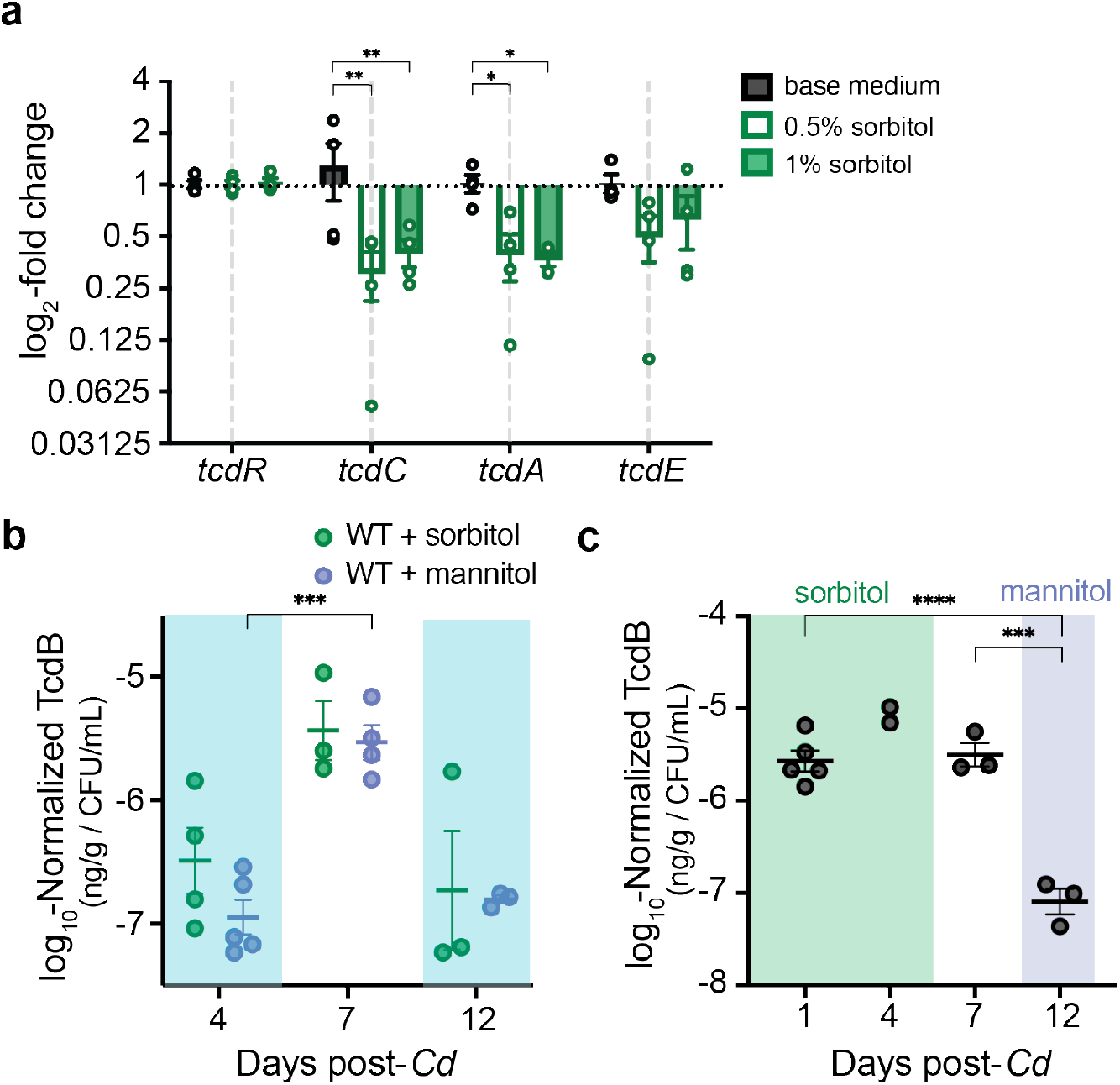
Excess sorbitol represses toxin production *in vitro* and *in vivo*. **a**, Minimal medium supplemented with 1% or 0.5% sorbitol leads to significantly lower expression of *tcdA* and *tcdE* after 8 hours growth compared to un-supplemented base medium (mean +/-SEM, n=4 biological replicates per condition. Two-way ANOVA across genes: F_(3,36)_=3.429, *P*=0.0271; across sorbitol supplementations: F_(2,36)_=11.17, *P*=0.0002 with Dunnett’s multiple comparisons test using base medium as control group for sorbitol supplementation comparisons within each gene). **b**, Presence of sorbitol or mannitol in drinking water leads to relatively lower toxin production *in vivo* (days 4 and 12, denoted by shaded boxes) compared to when sorbitol or mannitol are absent (day 7; mean +/-SEM, mixed effects analysis with Sidak’s multiple comparisons test: day is significant but group is not, F_(0.8915,7.132)_=18.37, *P* = 0.004). **c**, Addition of exogenous mannitol (day 12) leads to lower production of toxin *in vivo* in the *ΔsrlD* strain compared to sorbitol supplementation (days 1, 4) or regular water (day 7; mean +/-SEM, one-way ANOVA F_(2,8)_=45.18 with Tukey’s post-hoc multiple comparisons. Day 4 was excluded from the ANOVA, as only 2 data points are present). For all panels, * *P* < 0.05, ** *P* < 0.01, *** *P* < 0.001, **** *P* < 0.0001.

**Extended Data Fig. 5.**
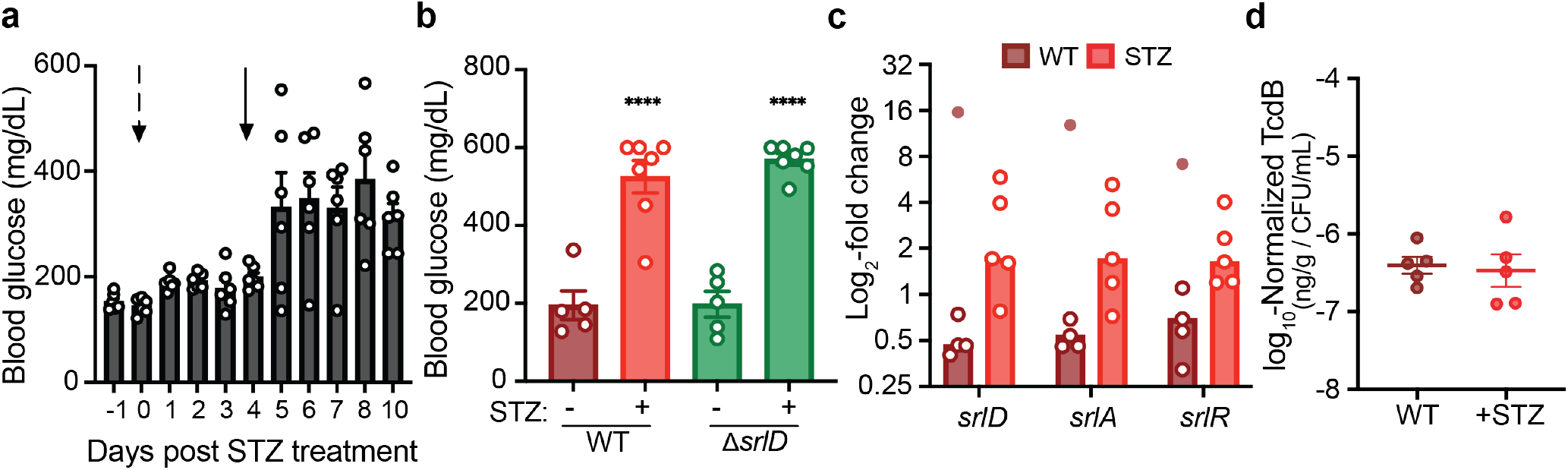
Streptozotocin treatment increases fasting blood glucose levels in conventional and mono-colonized mice. **a**, Development of streptozotocin (STZ)-induced hyperglycemia model in Swiss-Webster Excluded Flora mice. Mice were fasted for 4-6 hours prior to measurement of blood glucose levels via tail vein snip. An initial injection (Day 0, indicated by dashed arrow) of 4.5 mg STZ was insufficient to increase blood glucose levels. A larger dose of 9.1 mg STZ administered on day 4 (solid arrow) was sufficient to increase blood glucose (mean +/-SEM, n=6) and was used for subsequent experiments with *Cd* infection. **b**, Unfasted blood glucose levels of germ-free mice mono-colonized with WT or *ΔsrlD* at 3 days post-infection (mean +/-SEM, n=5-7 mice/group, **** *P* < 0.0001, one-way ANOVA with multiple comparisons; STZ-treated groups significantly different from non-treated). Mice were treated with STZ via IP injection 4 days prior to *Cd* infection. **c**, *Cd* expression of genes in the sorbitol utilization locus in conventional (WT) or streptozotocin-treated (STZ) mice. An outlier (Fig. 3c, tested for with robust nonlinear regression, Q=0.2%) from one RNA sample isolated from one mouse is indicated by the filled circle, bars denote median. **d**, Streptozotocin treatment does not alter toxin production *in vivo. Cd* Toxin B quantified in feces of conventional mice infected with WT *Cd* 24 hours post-infection.

**Extended Data Fig. 6.**
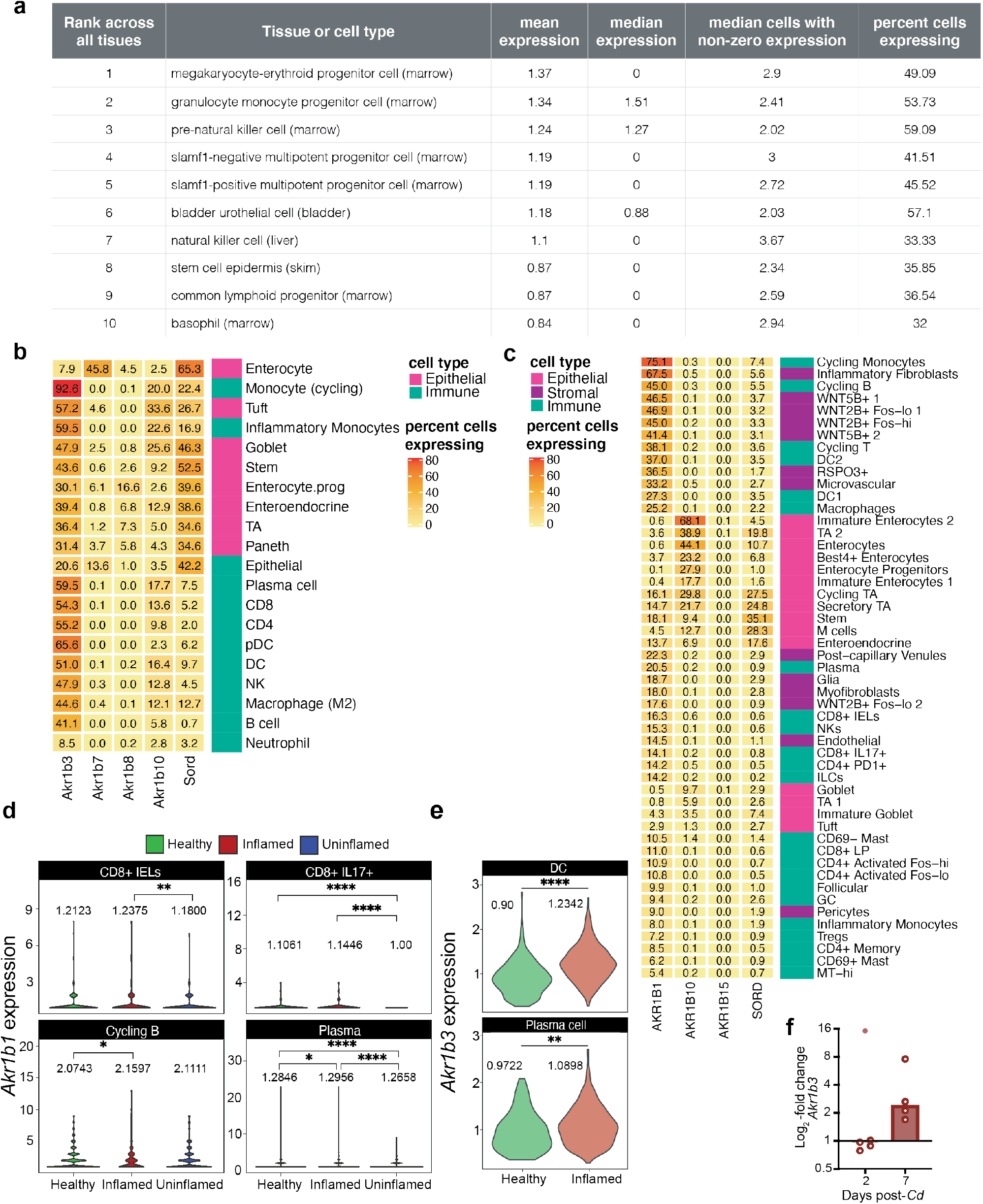
Aldose reductase is an immune cell-associated gene. **a**, Top 10 cell types with highest expression of *Akr1b3* across 20 mouse organs demonstrates high prevalence of AR in immune-associated cell types^35^. **b**, Percent of cells in mouse colonic tissue expressing isoforms of aldose reductase and sorbitol dehydrogenase. **c**, Percent of different cell types in human colonic explants expressing the three isoforms of aldose reductase and sorbitol dehydrogenase. **d**, *Akr1b1* expression (log_2_-TP10K+1) in cell types exhibiting significantly increased AR expression in inflammatory colonic explants from humans with ulcerative colitis (Inflamed) compared to within-subject non-inflamed tissue (Uninflamed) vs. healthy controls that do not have ulcerative colitis (Healthy; pairwise Wilcoxon-rank sum test across all immune cell types using non-zero expression levels. Means for each cell type are shown.). **e**, Dendritic cells (DC) and plasma cells in mouse large intestine exhibited significant increase in *Akr1b3* expression (log_2_-TPM+1) during infection with *H. polygyrus* (Wilcoxon-rank sum test across all immune cell types using non-zero expression levels. Means for each cell type shown.). **f**, Expression of *Akr1b3* in the proximal colon of conventional mice infected with WT *Cd*. Outlier (Fig. 2e, detection method: robust nonlinear regression, Q=0.2%) is denoted by the filled point, bars denote median.

**Extended Data Fig. 7.**
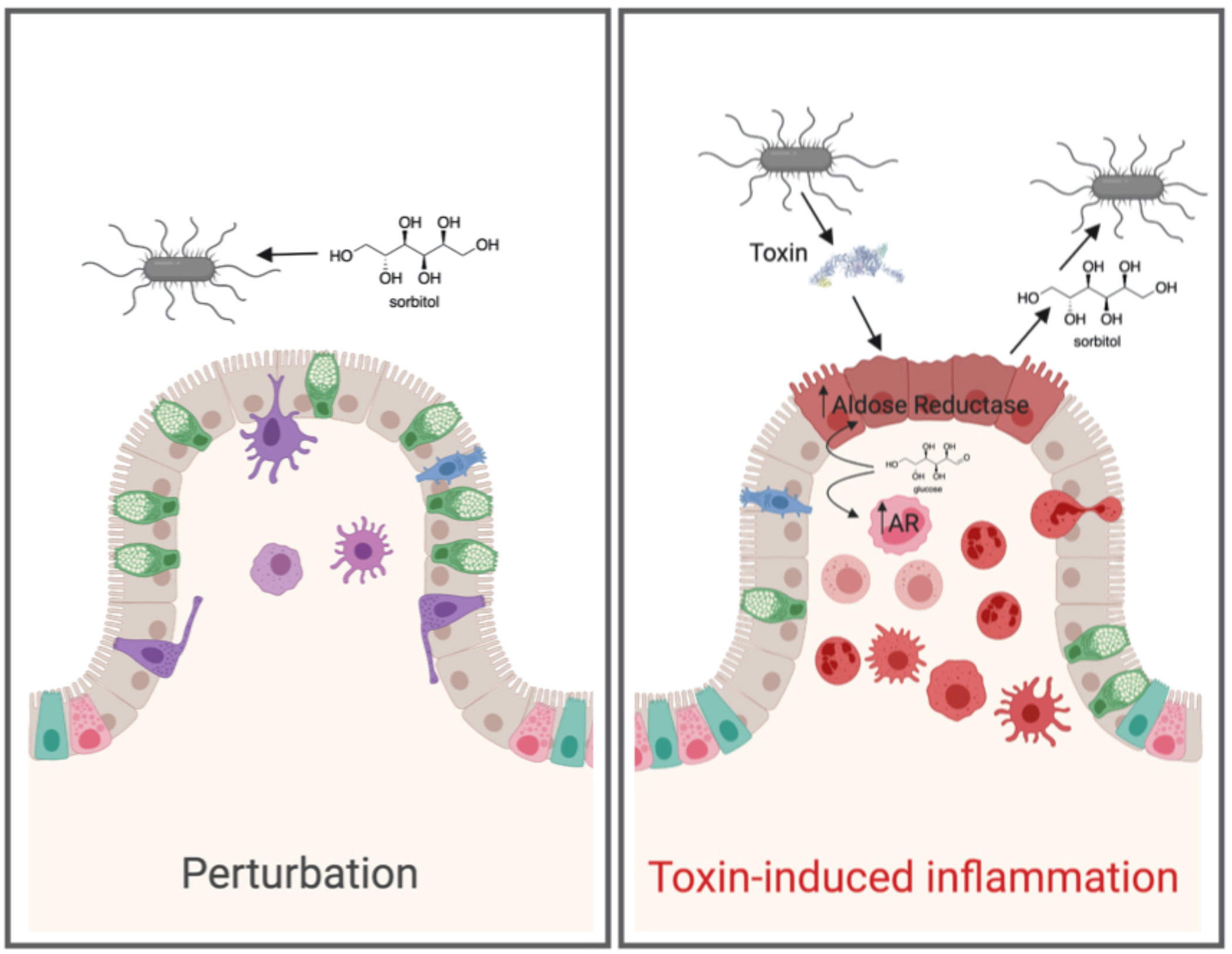
A model for *Cd* sorbitol utilization. *Cd* can utilize diet-derived sorbitol, which spikes after disturbance to the microbiota (left). Toxin-induced tissue damage (right) leads to up-regulation of host aldose reductase in the epithelium as well as recruitment of immune cells that express AR. *Cd* is able to utilize host-derived sorbitol.

**Extended Data Fig. 8.**
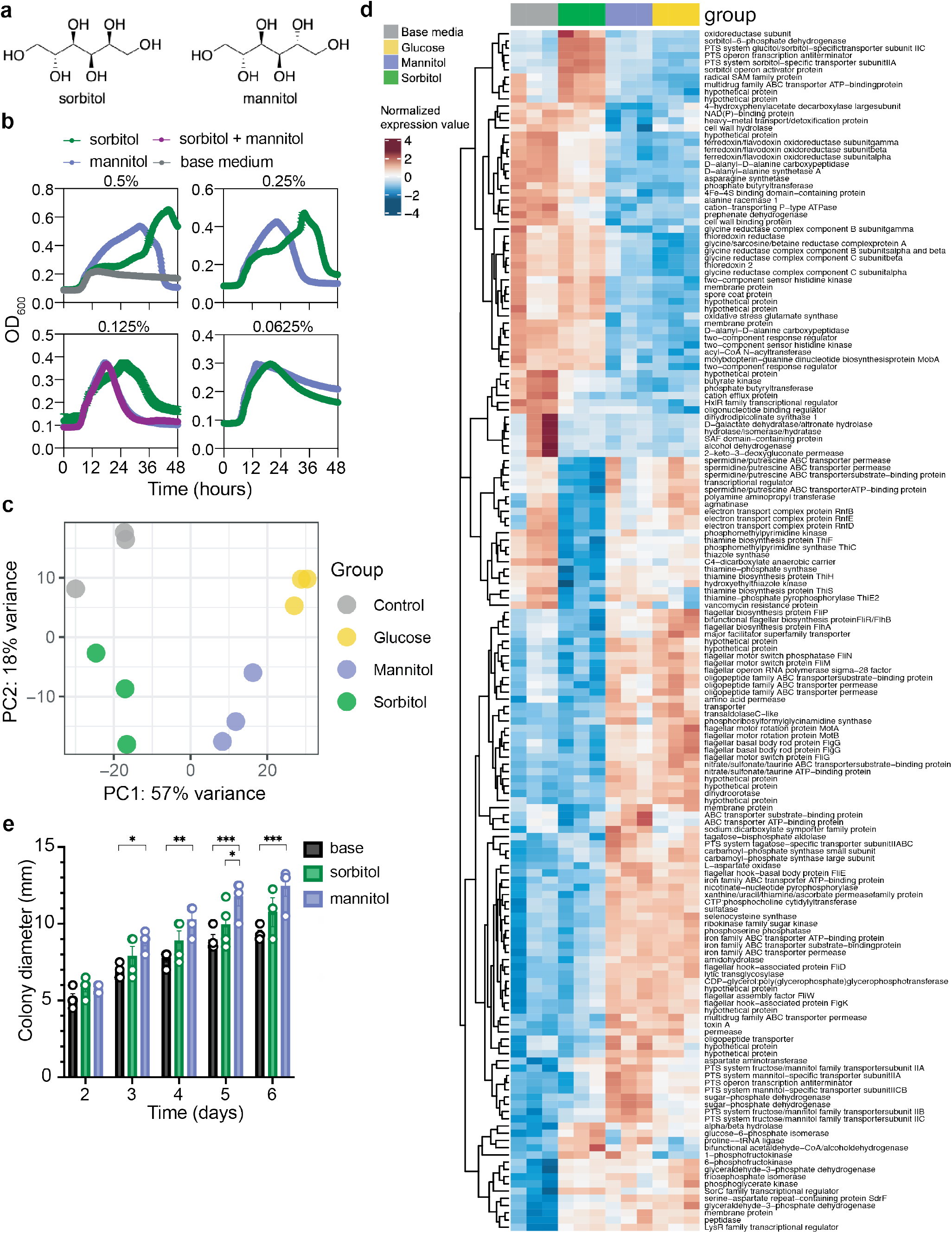
Sorbitol and mannitol lead to distinct metabolic programs *in vitro*. **a**, Chemical structures of isomers sorbitol and mannitol. **b**, Sorbitol and mannitol added to minimal medium engender distinct growth kinetics *in vitro* (mean +/-SEM, n=5 replicates for each growth condition). **c**, Principal components analysis of variance stabilizing-transformed RNA-seq count values from *Cd* grown for 11 hours in minimal medium (control, grey), or minimal medium supplemented with 0.25% sorbitol (green), mannitol (purple), or glucose (yellow). **d**, Significantly differentially expressed genes between sorbitol supplementation and base medium or mannitol supplementation (n=3 biological replicates; colors represent row-normalized variance stabilizing-transformed counts. Adjusted *P* value < 0.01, Wald test with Bonferroni p-adjust method.) **e**, Mannitol supplementation to 0.3% soft agar plates leads to significantly increased motility compared to base medium (days 3-6) and sorbitol supplementation (day 5). Sorbitol supplementation does not lead to a significant increase in motility compared to un-supplemented motility plates (* *P* < 0.05, ** *P* < 0.01, *** *P* < 0.001, repeated measures two-way ANOVA with Tukey’s multiple comparisons).

**Extended Data Fig. 9.**
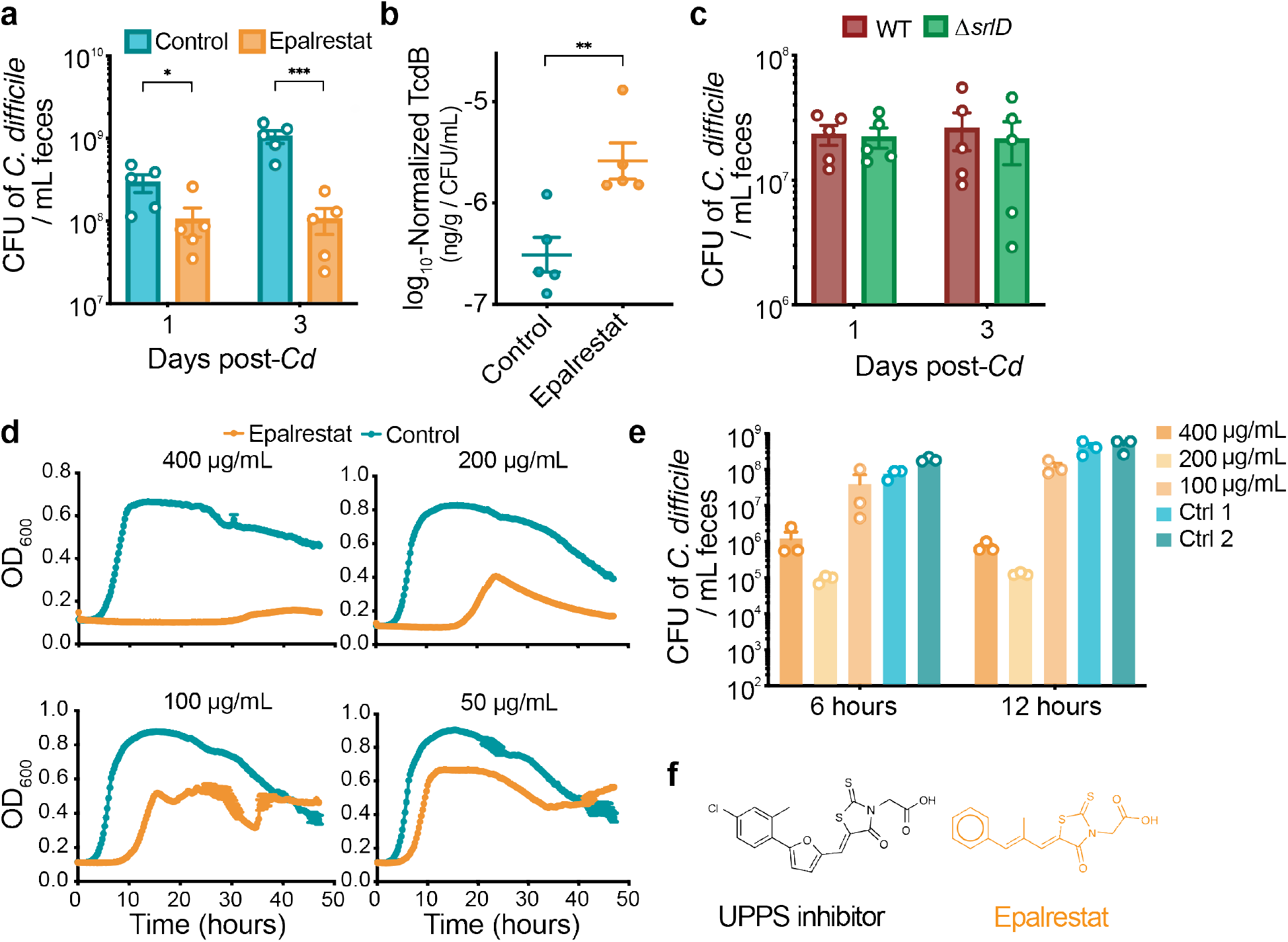
Epalrestat inhibits *Cd* growth *in vitro* and *in vivo*. **a**, Germ-free mice were mono-colonized with WT *Cd* and gavaged with the AR inhibitor epalrestat once daily or gavaged with vehicle control. Epalrestat treatment significantly reduces *Cd* abundance (mean +/-SEM, * *P* < 0.05, *** *P* < 0.001, unpaired t-test). **b**, In the presence of epalrestat, WT *Cd* produces relatively more toxin *in vivo* (median, ** *P* < 0.01, Mann-Whitney). **c**, The *ΔsrlD* mutant colonizes germ-free mice on standard diet equally well as WT *Cd* (mean +/-SEM). **d**, Epalrestat inhibits wild-type *Cd* 630ΔErm growth in rich medium in a dose-dependent manner (n=5 biological replicates). **e**, Absolute abundance of WT *Cd* after 6 or 12 hours of growth in rich medium is inhibited due to incubation with epalrestat (mean +/-SEM, n=3 biological replicates). **f**, Chemical structures of AR inhibitor epalrestat (bottom) and an antibiotic with activity against cis-prenyl transferase undecaprenyl diphosphate synthase (UPPS, top; ref. ^45^).

**Extended Data Table 1. Detailed results of blinded histopathological scoring for mice mono-colonized with WT or Tox-*Cd***.

**Extended Data Table 2. DESeq2 *in vivo* differential gene expression analysis from mice mono-associated with WT or Tox-*Cd***.

**Extended Data Table 3. *In vitro* differential gene expression analysis comparing different carbohydrate supplements to minimal medium**.

**Extended Data Table 4. List of primers used in this study**.

## Methods

### Mouse Strains and Husbandry

All animal experiments were performed in accordance with the Stanford Institutional Animal Care and Use Committee. Mice were maintained on a 12-hour light/dark cycle, fed *ad libitum*, and maintained in flexible film gnotobiotic isolators for the duration of all experiments (Class Biologically Clean, Madison WI). Swiss-Webster mice were utilized for gnotobiotic experiments and sterility of germ-free mice was verified by 16S PCR amplification and anaerobic culture of feces. Aldose reductase knockout mice (ARKO, *Akr1b3*^*-/-*46,47^) in C57/B6 background, C57/B6 WT mice (Jackson Laboratories), and Swiss-Webster Excluded Flora mice bred in-house were used for conventional experiments. All experiments were performed between 9-15 weeks of age and animals were sex and age-matched except for *in vivo* co-infections, for which sexes were mixed equally in all groups. Mice were fed an autoclaved standard diet (Purina LabDiet 5010 for conventional, 5K67 for gnotobiotic) in all experiments except where fully-defined diets lacking sorbitol were used: Figs. 2g,h, Extended Data Figs. 3,4b,c: BioServ Product #S6183, Fig. 4: Teklad Diet TD.86489 or BioServ #S6185. Defined diets were used starting 48 hours prior to *Cd* infection, and cage bedding was replaced immediately subsequent. For experiments in which sorbitol or mannitol (Sigma) were provided in drinking water, 1% (w/v) of the sugar alcohol was filter-sterilized and provided *ad libitum* starting immediately prior to *Cd* infection.

### Bacterial Strains and Growth Conditions

#### Cd culture conditions

Isogenic *Cd* toxin-deficient mutants and parental strains obtained from refs.^15,38^ harbored ClosTron insertional mutations in each of the toxin genes. *Cd* strains were cultured anaerobically (85% N_2_, 10% CO_2_, 5% H_2_) on BHIS plates: Brain Heart Infusion Agar (BD Difco) supplemented with 0.5 mg/mL cysteine and 5 mg/mL yeast extract. Liquid medium used was Brain Heart Infusion broth (BD Difco) supplemented with 5 mg/mL yeast extract and 1 mg/mL cysteine. Minimal medium for growth curves, *in vitro* competition experiments, and *in vitro* RNA-seq was *C. difficile* defined medium (CDDM^48^) with no glucose added.

#### Selective culturing of C

difficile *from stool samples:* Feces were collected from individual mice, 2 biological replicates of 1 µl feces resuspended in 200 µl 1X sterile PBS, serially diluted 1:10, and 10 µl of each dilution plated on CDDC selective medium (*Clostridium difficile* agar base (Oxoid) supplemented with 7% defibrinated horse blood, 0.5 mg/mL cysteine, 250 µg/mL D-cycloserine, and 16 µg/mL cefoxitin). Colony-forming units (CFUs) were counted after 24 hours anaerobic growth at 37 °C.

#### Culture conditions for defined community members

*Bacteroides thetaiotaomicron* VPI-5482, *Bacteroides fragilis* NCTC 9343, *Clostridium sporogenes* ATCC 15579, *Clostridium symbiosum* BEI #HM-309, and *Bifidobacterium infantis* ATCC 15697 were cultured anaerobically. *B. thetaiotaomicron* and *B. fragilis* were cultured on BHI Blood Agar plates (Brain Heart Infusion supplemented with 10% defibrinated horse blood and 200 µg/mL gentamycin) or BHIS for Bacteroides (Brain Heart Infusion supplemented with 5 µg/mL hemin and 2 µg/mL vitamin K1), which was also used for liquid medium. *C. sporogenes* was cultured in TYG, both solid and liquid medium (30 g/L tryptone, 20 g/L yeast extract, 1 g/L sodium thioglycolate; plates were supplemented with 125 µg/mL D-cycloserine, 38 µg/mL sulphamethoxazole, and 2 µg/mL trimethoprim). *Escherichia coli* K-12 MG1655, *Edwardsiella tarda* ATCC 23685, and *Proteus penneri* ATCC 35198, were cultured aerobically in LB medium supplemented with 10 µg/mL vancomycin or BHIS for Bacteroides. *B. infantis* and *C. symbiosum* were cultured in MRS (broth or agar, Sigma) and BHIS for Bacteroides.

### C. difficile infection

For all mouse experiments, animals were gavaged with 200 µl culture *Cd* grown in 5 mL BHIS broth for 16 hours, corresponding to ∼5×10^7^-1×10^8^ CFU/mL. Cultures were prepared for gavage anaerobically in sterile 2 mL cryovials with inner threading. For co-infections, a 1:1 mixture of each strain was used. No antibiotic pre-treatment was used for gnotobiotic experiments (mono-colonizations or defined community experiments). For conventional experiments (SWRF), mice were each gavaged with 1 mg clindamycin filter-sterilized in 200 µl water 24 hours prior to infection. Bedding was replaced immediately after both antibiotic administration and *Cd* gavage.

For weight loss comparisons between wild-type B6J and ARKO mice, mice were administered an antibiotic cocktail *ad libitum* in water for 3 days (0.4 g kanamycin, 0.035 g/L gentamycin, 0.057 g/L colistin, 0.215 g/L metronidazole, 0.045 g/L vancomycin^42^), ending 2 days prior to clindamycin treatment. Bedding was replaced at the onset of antibiotic cocktail administration and then mixed between ARKO and WT groups upon removal of antibiotic water to mitigate potential host genotype-specific influences on microbiota composition. In accordance with other conventional mouse experiments, clean bedding was replaced at the time of clindamycin treatment as well as *Cd* gavage.

For all data shown, mice were infected with the parental strain *C. difficile* 630ΔErm (WT) or mutant derivatives thereof (Tox-or *ΔsrlD*), except for tissue qRT-PCR data presented in Figure 3d, for which mice were infected with the wild-type strain *C. difficile* R20291 or its triple-toxin isogenic mutant, TcdA^-^B^-^ CDT^-38^.

#### Defined community associations

For the relative abundance (Extended Data Fig. 1a) data and host polyol pathway expression data (Fig. 3d), germ-free mice were tri-colonized with *B. thetaiotaomicron, E. coli*, and *C. sporogenes*. Separate overnight cultures for each strain were combined anaerobically in equal parts and administered via 200 µl gavage. Commensal members equilibrated for ten to thirteen days prior to infection or gavage with anaerobic PBS for the uninfected control. For host expression data, mice were infected with WT R20291 or its triple-toxin isogenic knockout, TcdA^-^B^-^CDT^-38^

For gnotobiotic defined community experiments in Figs. 2g,h, and Extended data Figs. 3,4, overnight cultures of *B. fragilis, B. infantis, C. symbiosum, P. penneri*, and *E. tarda* were combined and administered to germ-free mice via gavage in 200 µl. Communities were allowed to stabilize on standard diet for 13 days prior to switching to the fully-defined diet 48 hours before *Cd* infection.

### Generation of *Cd* mutant

The *C. difficile* 630ΔErm*ΔsrlD* mutant was constructed via the *pyrE* allelic exchange system as previously described^49^. Briefly, regions 1kb up- and down-stream of the targeted deletion region were amplified from genomic DNA and inserted into the pMTL-YN3 vector backbone via Gibson Assembly, transformed into and propagated in *Escherichia coli* TG1 prior to transformation into *E. coli* HB101/pRK24 conjugation-proficient cells. The pMTL-YN3 deletion plasmid was transferred to *C. difficile* 630ΔErmΔpyrE via spot-plate mating conjugation. During conjugation and proximal selection steps, *Cd* and *E. coli* were cultured anaerobically with higher atmospheric H_2_ atmospheric levels (85% N_2_, 5% CO_2_, 10% H_2_). Plasmid integrants were selected on BHIS medium supplemented with 15 µg/mL thiamphenicol, 50 µg/mL kanamycin, 16 µg/mL cefoxitin, 5 µg/mL uracil, and deletion mutants selected for on CDDM supplemented with 5 µg/mL uracil and 2 mg/mL 5-fluoroorotic acid (5-FOA). Deletion loci were sequenced verified, after which the mutant *pyrE* locus was restored with a second round of mutagenesis using the pMTL-YN1C plasmid.

### Histopathology scoring

Tissue segments of approximately 1 cm in length were taken from the cecal blind tip and proximal colon of mice, fixed in Formalin for 24-48 hours and then transferred to 70% ethanol for long-term storage. Tissues were embedded in paraffin, sectioned and stained with haematoxylin and eosin (H&E). Two sections from each tissue were given scores of 0-3 for each of the following parameters: inflammatory cell infiltration, mucosal hyperplasia, vascular congestion, epithelial disruption, and submucosal edema. The cumulative lesion score is the sum of each score in these independent categories. Scoring was performed blinded.

### 16S sequencing and analysis

Total DNA was extracted from frozen fecal samples using the DNeasy PowerSoil HTP 96 kit (Qiagen). Barcoded primers were used to amplify the V3-V4 region of the 16S rRNA gene using 515f and 806r primers, as described previously^50,51^. Amplicon clean-up was performed using the Ultra Clean 96 well PCR Clean Up kit (Qiagen) prior to fluorescent quantification of DNA yield (Quant-iT), library pooling, and quality control on BioAnalyzer. 300 bp paired-end reads were generated on Illumina MiSeq2500. Demultiplexing was performed with ’split_libraries_fastq.py’ in QIIME 1.9.1^52^ and reads were assigned to a custom 16S rRNA reference database of the defined community using ’pick_closed_reference_otus.py’.

### RNA-sequencing and analysis

#### RNA isolation

RNA from cecal contents was isolated using the Qiagen RNA Power Microbiome kit per the manufacturer’s instructions. RNA from culture was extracted with the Qiagen RNEasy Mini kit. 10 mL culture was spun at 5,000 x *g* for 10 minutes, resuspended in kit buffer RLT + ßME, and bead-beat using acid-washed glass beads for 5 minutes at 4°C.

#### Library prep and Sequencing

Ribosomal RNA was depleted from total RNA isolated from cecal contents using the Illumina Ribo-Zero Gold Epidemiology rRNA Removal Kit. No experimental rRNA removal was used for the *in vitro* RNA-seq experiment. Depletion of rRNA as well as RNA quality was verified with Agilent Bioanalyzer Prokaryote Total RNA Pico prior to moving forward with library preparation using the Illumina TruSeq mRNA Stranded HT Library Prep kit. Sequencing for the *in vivo* RNA-seq was performed on an Illumina HiSeq4000 instrument with 100 bp paired-end reads. *In vitro* RNA-seq was performed on Illumina NovaSeq with 150 bp reads.

#### Analysis

Raw, paired reads were imported into CLC Genomics Workbench version 11 with a maximum distance of 1000 bp. *In vivo* reads were trimmed with a quality limit of 0.05, an ambiguous limit of 2, based on automatic read-through adaptor trimming, with a minimum number of nucleotides of 50. Broken pairs and discarded sequences were not saved. Reads were mapped to the *C. difficile* 630 genome (RefSeq Accession NC_009089.1) using standard parameters: paired reads were mapped to the gene track with a mismatch cost of 2, insertion cost of 3, deletion cost of 3, length and similarity fractions of 0.8. Read counts for each gene were exported and remaining ribosomal RNA reads were manually removed prior to differential expression analysis in R with DESeq2^53^.

Genomic loci with fewer than 3 reads across all samples were filtered and removed. Standard parameters within DESeq2 were used for differential expression analysis: pairwise comparisons across all groups using the Wald test with a significance threshold of *P* < 0.01 was used to identify significantly differentially expressed genes. Row-normalized variance stabilization-transformed (*vst*) counts are presented in the heatmap for Extended Data Fig. 8d.

Pathway enrichment analysis for the *in vivo* RNA-seq was conducted using ShinyGO^54^ with the *C. difficile* String-DB reference using the UniProt code for genes significantly up-regulated in either WT or Tox-conditions. Six genomic loci annotated as pseudogenes were not included in the analysis.

### Toxin B quantification

Free toxin concentration in feces or cecal contents was quantified with the tgcBIOMICS Toxin B ELISA kit. 25-50 mg frozen fecal sample was thawed, weighed, and resuspended in 450 µl dilution buffer prior to proceeding via the manufacturer’s instructions. Toxin levels were quantified (ng toxin per g feces) using an external standard curve and normalized to the absolute abundance of *Cd* levels (CFU / mL) as previously described^44^.

### qRT-PCR

Total RNA from feces or culture was extracted as described above. RNA from frozen host proximal colon tissue was extracted with the RNEasy Mini Kit: 25-30 mg tissue was bead-beat with acid-washed glass beads in 600 µl RLT + ßME for 5 min at 4°C prior to proceeding with the kit protocol. cDNA was generated using random primers and Superscript III Reverse Transcriptase per the manufacturer’s instructions; RNase OUT was included. QPCR was performed with SYBR Brilliant III on an Agilent MX3000P instrument in 20 µl volumes with 0.3 µl reference dye added. Changes to expression were calculated by normalizing Ct value to a housekeeping gene and a control condition with the delta-delta Ct method.

### Competition assays

*In vivo* and *in vitro* competition between wild-type and *ΔsrlD Cd* was assessed using qPCR. Primer pairs (listed in **Extended Data Table 4**) targeting the wild-type *srlD* locus (WT), surrounding the *srlD* deletion (*ΔsrlD*, used for *in vivo* competition experiments), or the *LacZ* locus, which is complemented in tandem with the restored *pyrE* gene using the pMTL-YN1C plasmid (*ΔsrlD*, used for *in vitro* competition experiments) were validated with eight 1:4 serial dilutions of purified genomic DNA from WT or *ΔsrlD Cd*. QPCR reactions were performed as described above. For *in vitro* competition assays, the ratio of each genotype was calculated relative to the *Cd* housekeeping gene *rpoA*, normalizing to levels with 0% sorbitol. For *in vivo* competition assays, the efficiency value (*E*) for each primer pair was calculated as 10^(1/-slope)^ of log_10_(DNA input) against Ct value. The competitive index of each genotype was calculated as follows: *E*^-Ct^ WT primer pair / *E*^-Ct^ *ΔsrlD* primer pair. A competitive index of 1 indicates that each strain is at equal abundance.

### Streptozotocin-induced hyperglycemia

A single high-dose injection of Streptozotocin was used to induce hyperglycemia in mice. Mice were fasted for 4-6 hours prior to IP injection of 350 µl 26 mg/mL Streptozotocin prepared in sterile sodium citrate buffer pH 4.5. Conventional mice were fasted for 4-6 hours prior to blood glucose measurements via tail vein snip with OneTouch blood glucose monitor. Blood glucose measurements taken in gnotobiotic mice at experimental endpoint were not fasted.

### Sorbitol and mannitol quantification

A GCMS-based mass spectrometry assay was developed to quantify mannitol and sorbitol from biological samples. Fecal samples were dried at 70°C for 1-1.5 hours, weighed, and metabolites were extracted by bead-beating in 500 µl LCMS-grade MeOH. Samples were centrifuged at 16,000 x *g* for 10 minutes. 200 µl supernatant was mixed thoroughly with 400 µl dichloromethane and d_8_-sorbitol was added as an internal standard. Samples were evaporated completely at 70°C under high-purity nitrogen, and then derivatized with trimethylsilyl reagents for 1 hour at 70°C using a 8:2:1 ratio of pyridine:hexamethyldisilazane:chlorotrimethylsilane. Samples were centrifuged at 16,000 x g for 10 minutes; supernatant was transferred to autosampler vials and 1 µl injected into an Agilent 7890A GC System equipped with 5975C inert MSD with Triple-Axis Detector using electron ionization (EI). A J&W DB-5MS UI column (30 m length, 0.250 mm ID, 0.25 film thickness; Agilent PN 122-5332UI) was run at 1 mL/min helium, with the following temperature program: 180°C, 1 minute initial hold; ramp 1 5°C/min to 215°C; ramp 2 40°C/min to 320°C, 5 minute hold. Ion chromatogram extraction and peak integration were performed using Agilent ChemStation Enhanced Data Analysis software. Product ions used to identify mannitol and sorbitol: m/z 205, 217, and 319. Product ions for d_8_-sorbitol: m/z 208, 220, and 323. Sorbitol and Mannitol standards were derivatized as described above and used to generate standard curves to enable sugar alcohol quantification.

### Analysis of published datasets

Tissue and cell-specific expression of AR from *Tabula muris* (https://tabula-muris.ds.czbiohub.org/)^35^ was searched for *Akr1b3*. Two single-cell sequencing studies targeting the large intestine^36,37^ were searched for all isoforms of AR and sorbitol dehydrogenase using the Single Cell Portal (https://singlecell.broadinstitute.org/single_cell). The tissue type (immune, epithelial, stromal) was designated using metadata provided by each study. For AR expression response to inflammatory conditions, data was downloaded and cells with non-zero expression levels were included in the analysis. Pairwise Wilcoxon rank-sum tests (implemented with *rstatix*) were conducted across all immune cell types; those with significantly increased AR expression in inflammatory conditions are shown.

AR expression in response to purified *Cd* TcdA injection in the cecum^39^ was compared to the sham control at 16 hours and was implemented in GEO2R (NCBI, GSE44091). We performed a differential correlation analysis^55^ to identify genes positively correlated with AR in response to TcdA. The three microarray probes annotated as *Akr1b3* were used as input for the differential correlation analysis. An adjusted *P*-value cut-off of *P* < 0.05 was used to identify genes that positively correlate in response to TcdA and negatively or do not correlate in control mice. This gene set was used to conduct gene set enrichment analysis with Bonferroni multiple hypothesis correction using Panther (Panther.org) with the Reactome database.^56^

### Statistical Analysis

All statistics were performed in R (R Core Team, 2018) and Prism (GraphPad), and visualized with ggplot2^58^ and Prism.

### Supplementary Text

#### Sorbitol and its stereo-isomer mannitol lead to unique transcriptional responses *in vitro*

As sorbitol and mannitol are both components of diet but the sorbitol utilization operon was uniquely up-regulated in inflammatory conditions, we compared the influence of these sugar alcohols on *Cd* physiology. Although the two sugars are stereoisomers (**Extended Data Fig. 8a**), *Cd* exhibited different growth kinetics when minimal medium was supplemented with mannitol or sorbitol (**Extended Data Fig. 8b**). Our data confirm previous descriptions of sorbitol utilization by *Cd* as a secondary carbon source, consumed after other rapidly-metabolizable carbohydrates, including mannitol^31^.

We explored whether transcriptional profiles during *in vitro* growth could provide insight into how sorbitol and mannitol differentially impact *Cd*. Carbohydrate-free minimal medium was supplemented with sorbitol, mannitol, or glucose, and total RNA was extracted for transcriptional profiling, using un-supplemented medium as a reference. The three sugars led to distinct transcriptional profiles (**Extended data Fig. 8c**): sorbitol supplementation elicited 65 differentially expressed genes compared to base medium, 127 genes were differentially expressed between sorbitol and mannitol, 248 between sorbitol and glucose, but only 54 between glucose and mannitol (Wald tests with Bonferroni correction, adjusted *P* value < 0.01, **Extended Data Table 3**).

Visualizing expression of genes that were significantly different between sorbitol supplementation and mannitol or base medium (**Extended Data Fig. 8d**) highlighted the common metabolic programs induced by mannitol and glucose, which were distinct from genes induced by sorbitol. A set of metabolic genes were specifically down-regulated with exogenous sorbitol addition compared to the other three conditions, including the rnf electron transport complex, thiamine (vitamin B1) biosynthesis, and polyamine metabolism. Mannitol and glucose supplementation corresponded to increased expression of several metabolic programs, as well as motility, flagellar biosynthesis, and chemotaxis (**Extended Data Fig. 8d**). Accordingly, mannitol supplementation led to a significant increase in WT *Cd* motility in soft agar motility plates, as opposed to sorbitol supplementation, which was not different from base medium (**Extended Data Fig. 8e**). Our transcriptional profiling confirms previously described differential regulation of mannitol and sorbitol metabolism by *Cd*^31^, and with measurable motility differences, we identify a phenotypic correlate of these nutrient conditions.

### Aldose reductase inhibitor epalrestat inhibits *Cd in vivo* and *in vitro*

To decrease host aldose reductase activity in gnotobiotic mice, we employed the AR inhibitor epalrestat^30,59^. When mice were gavaged with epalrestat, we observed a decrease in *Cd* abundance in mono-colonized mice (**Extended Data Fig. 9a**) and an increase in relative TcdB production (**Extended Data Fig. 9b**). However, the *ΔsrlD* mutant was not attenuated in colonization in mono-colonized mice (**Extended Data Fig. 9c**). To distinguish whether the effect of epalrestat on *Cd* abundance in mono-colonization was due to an altered host immune response^60^ or a direct effect of epalrestat on *Cd*, we cultured wild-type 630ΔErm in rich medium in the presence of epalrestat and observed a dose-dependent inhibition of *Cd* growth *in vitro* as measured by optical density (**Extended Data Fig. 9d**) and CFUs (**Extended Data Fig. 9e**). It was previously noted in a computational screen for novel antibiotics targeting the *cis*-prenyl transferase undecaprenyl diphosphate synthase (UPPS), essential for peptidoglycan biosynthesis, that a top target against MRSA, *Listeria monocytogenes, Bacillus anthracis*, and a vancomycin-resistant *Enterococcus spp*. was strikingly similar in structure to epalrestat (**Extended Data Fig. 9f**, ref. ^45^). Thus, it is likely that epalrestat inhibits *Cd* cell wall biosynthesis to limit growth *in vivo*.

## Data Availability

Raw RNA-seq source data is available through the NCBI Sequence Read Archive (SRA). *In vivo* RNA-seq (Extended Data Table 2): PRJNA666929, *in vitro* RNA-seq (Extended Data Table 3): PRJNA667108. Code is available upon request.

## Author contributions

K.M.P. and J.L.S. conceived the project idea, designed experiments, and wrote the manuscript. K.M.P. executed experiments and performed data analysis.

## Acknowledgements

The authors would like to give special thanks to Steven Higginbottom for assistance with all animal experiments. Allis Chien at the Stanford University Mass Spectrometry Facility provided assistance with GC-MS protocol development. Sarah Kuehne and Nigel Minton from the University of Nottingham provided the toxin-mutant *C. difficile* strains and Aimee Shen provided reagents for generation of new *C. difficile* mutants. Aruni Bhatnagar and Donald Mosely from the University of Louisville generously provided the aldose reductase knockout mice. Dawn Davis shared protocols and advice regarding the development of the streptozotocin model of hyperglycemia. BioRender was used to generate Extended Data Figs. 1b and 7. This study was supported by R01-DK08502510 (special thanks to Bob Karp for his service and support at NIDDK) to J.L.S. and a Ford Foundation Fellowship and NSF Graduate Fellowship to K.M.P. The authors thank all members of the Sonnenburg laboratory who provided thoughtful feedback throughout the development of the project.

